# RBM45 associates with nuclear stress bodies and forms nuclear inclusions during chronic cellular stress and in neurodegenerative diseases

**DOI:** 10.1101/856880

**Authors:** Mahlon Collins, Yang Li, Robert Bowser

**Affiliations:** Departments of Neurobiology, Barrow Neurological Institute, Phoenix, AZ, USA; Neurology, Barrow Neurological Institute, Phoenix, AZ, USA; Department of Neurobiology, University of Pittsburgh, Pittsburgh, PA, USA

## Abstract

RBM45 is a multifunctional RNA binding protein (RBP) found in cytoplasmic and nuclear inclusions in amyotrophic lateral sclerosis (ALS), frontotemporal lobar degeneration (FTLD), and Alzheimer’s disease (AD). While cytoplasmic RBM45 inclusions contain other disease-associated proteins, nuclear RBM45 inclusions are morphologically and biochemically distinct from previously described nuclear inclusion pathology in these diseases. To better understand nuclear RBM45 aggregation and inclusion formation, we evaluated the association of RBM45 with a variety of membraneless nuclear organelles, including nuclear speckles, Cajal bodies, and nuclear gems. Under basal conditions, RBM45 is diffusely distributed throughout the nucleus and does not localize to a specific nuclear organelle. During cellular stress, however, the nuclear RBM45 distribution undergoes an RNA-binding dependent rearrangement wherein RBM45 coalesces into a small number of nuclear puncta. These puncta contain the nuclear stress body (NSB) markers heat shock factor 1 (HSF1) and scaffold attachment factor B (SAFB). During chronic stress, the persistent association of RBM45 with NSBs leads to the formation of large, insoluble nuclear RBM45 inclusions. RBM45 nuclear inclusions persist after stressor removal and NSB disassembly and the inclusions resemble the nuclear RBM45 pathology seen in ALS, FTLD, and AD. We also quantified the cell type- and disease-specific patterns of RBM45 pathology in ALS, FTLD, AD, and non-neurologic disease control subjects. RBM45 nuclear and cytoplasmic inclusions are found in neurons and glia in ALS, FTLD, and AD but not in controls. Across diseases, RBM45 nuclear inclusion pathology occurs more frequently than cytoplasmic RBM45 inclusion pathology and exhibits cell type-specific variation. Collectively, our results define new stress-associated functions of RBM45, a mechanism for its nuclear aggregation and inclusion formation, a role for NSBs in the pathogenesis of diseases such as ALS, FTLD, and AD, and further underscore the importance of self-association to both the normal and pathological functions of RBPs in these diseases.

## Introduction

RBM45 is a developmentally regulated RNA binding protein (RBP) that forms inclusions in neurons and glia in amyotrophic lateral sclerosis (ALS), frontotemporal lobar degeneration (FTLD), and Alzheimer’s disease (AD)^1, 2^. Inclusions containing RBM45 occur in two distinct forms. Cytoplasmic RBM45 inclusions are positive for both TDP-43 and ubiquitin, and exhibit a skein-like or globular morphology. In contrast, nuclear RBM45 inclusions are punctate in appearance, negative for TDP-43 and ubiquitin, and have not been linked to a specific nuclear organelle or structure^1^. By virtue of their absence in non-neurologic disease controls, nuclear RBM45 inclusions likely result from one or more pathological processes. The mechanisms leading to their formation, the impact of nuclear RBM45 inclusions on cellular viability, and the cell-type specific patterns of nuclear RBM45 pathology across diseases remain unknown, however.

To better understand the normal functions RBM45 and how these relate to inclusion formation, we previously characterized functional protein domains in the RBM45 protein. By virtue of a bipartite nuclear localization sequence (NLS), RBM45 is a predominantly nuclear protein under basal conditions^3–5^. Mutation of the RBM45 NLS sequence leads to cytoplasmic sequestration of the protein, with attendant incorporation into cytoplasmic stress granules (SGs)^3, 4^. RBM45 also contains 3 RNA recognition motifs (RRMs) that govern its binding to RNA sequences^3, 4, 6^. Finally, RBM45 contains a novel “homo-oligomer assembly” (HOA) domain, an intrinsically disordered peptide sequence near the center of the protein (amino acids 258-318 of 474) that mediates the ability of RBM45 to self-associate and interact with other RBPs (including TDP-43 and FUS)^3, 6, 7^. These results demonstrated that RBM45 is capable of assembling into high-molecular weight oligomers, presumably as part of its normal function(s)^3, 7^. This type of RBP self-association or oligomerization is often a critical determinant of RBP incorporation into membraneless organelles such as SGs, P-bodies, nucleoli, paraspeckles, and Cajal bodies^8, 9^. We then used immunoprecipitation coupled to mass-spectrometry (IP-MS) experiments to identify RBM45-interacting proteins and infer putative biological functions of the protein. These results suggested that RBM45 regulates mRNA splicing and spliceosome function in the nucleus^5^. RBM45 may therefore be a component of one or more nuclear organelles that regulate mRNA splicing and processing, such as nuclear speckles or Cajal bodies.

The self-association and incorporation of RBPs in membraneless organelles also contributes to their pathological aggregation in diseases such as ALS and FTLD^8, 10–13^. A well-studied example of this phenomenon is the RBP aggregation that results from RBP incorporation and prolonged confinement in cytoplasmic SGs^14, 15^. SGs are cytoplasmic protein-mRNA complexes that form rapidly in response to cellular stress^16, 17^. They protect mRNAs from a harmful cellular environment and stall their translation, thus conserving energy and prioritizing translation of stress response-associated proteins (reviewed in ^16, 17^). TDP-43 and FUS are components of SGs. Chronic entrapment of these proteins in SGs results in the formation of insoluble inclusions containing these proteins in a variety of model systems^15, 18–21^. Likewise, SG marker proteins are also found in TDP-43 and FUS-positive neuronal cytoplasmic inclusions in ALS and FTLD patients^21, 22^,. The presence of low complexity and intrinsically disordered domains in these and other SG-associated RBPs facilitates aggregation and can lead to the formation of several forms of amyloid or amyloid-like structures, including amyloidogenic oligomers and fibrils^23–26^. This pathological “maturation” of RBP assemblies into insoluble aggregates and inclusions removes RBPs from their normal cellular milieu, leading to loss of normal RBP function(s)^27–30^. We propose that an analogous process occurring in the nucleus could contribute to both the normal functions of RBM45 and to the formation of the RBM45 nuclear inclusions seen in ALS, FTLD, and AD.

To test this hypothesis, we explored the association of RBM45 with a variety of membraneless nuclear organelles, including nuclear speckles, Cajal bodies, nuclear gems, and nuclear stress bodies (NSBs). We found that RBM45 is recruited to NSBs in human cell lines following exposure to a diverse array of cellular stressors. This association is dependent on the protein’s NLS and ability to bind RNA. The persistent association of RBM45 with NSBs is sufficient to promote its aggregation into insoluble complexes and, unlike NSBs, these complexes do not disassemble, but instead increase in size and number in the nucleus. Collectively, our results (1) describe novel functions of RBM45, (2) define a new mechanism by which the stress-associated functions of RBPs contribute to their aggregation and inclusion formation, and (3) provide quantitative measures of the cell type and disease-specific patterns of RBM45 pathology that occur in ALS, FTLD, and AD.

## Methods

### Cell Culture, Transfection, and Nuclear Stress Body Induction

HEK293 cells or FreeStyle 293F cells (Invitrogen, Waltham, MA, USA) were cultured in DMEM medium with 10% FBS and 1% Pen-Strep at 37°C with 5% CO_2_. Transfection was performed using the Lipofectamine 3000 reagent (Invitrogen) according to the manufacturer’s protocol and transfected cells were harvested 48 or 72 hours post-transfection. Tet-inducible EGFP-RBM45 stable cell lines (see below) were maintained in the presence of 50μg/ml Zeocin (Invivogen, San Diego, CA, USA). The expression of EGFP-RBM45 was induced with 1 μg/ml doxycycline hyclate (Sigma, St. Louis, MO, USA) for 16 hours before live-cell imaging.

To induce nuclear stress body (NSB) formation, we used a variety of previously described stressor protocols. Heat shock was performed by incubating the cells in a 42°C environment for various time lengths (ranging from 0.5 to 2 hours, as indicated in the text/figures) followed by various recovery times, ranging from 1 to 24 hours, as indicated^31, 32^. The following reagents were added to culture medium at the indicated concentrations to induce NSB formation, as previously described: 5-30 µM CDSO_4_ (2-24 hours)^31, 33^, 200-400 µM H_2_O_2_ (2-24 hours)^31, 34^, 0.1-1 mM sodium arsenite (0.25-24 hours)^35^, or 0.5-20 µM mitoxantrone (MTX; 2-24 hours)^36^.

### Plasmid Construction

For expression of WT and domain deletion RBM45 constructs, a plasmid containing RBM45, cGST-hRBM45 (HsCD00356971), was obtained from the DNASU Plasmid Repository at Arizona State University, Tempe, AZ, USA. The RBM45 cDNA was amplified by PCR using Phusion High-Fidelity DNA Polymerase (NEB) and sub-cloned into the pcDNA3 vector (Invitrogen). A 3xFLAG tag (DYKDHDGDYKDHDIDYKDDDDK) or 2xHA tag (DYPYDVPDYAGGAAYPYDVPDYA) was appended to the N-terminus of RBM45 to generate the 3xFLAG- or 2xHA-tagged constructs, respectively. In previous experiments, we found that appending tags to the C-terminus of the protein alters its subcellular localization, presumably by interfering with the function of the C-terminal nuclear localization sequence (amino acids 454-472 of 474). Thus, all RBM45 tags were appended to the protein’s N-terminus. Domain deletions and mutations were introduced by site-directed mutagenesis using overlap extension PCR^37^. Sequences of RBM45 domain deletion and non-functional NLS mutant constructs were as in Li et al (2015)^3^. For live-cell imaging of RBM45 dynamics during conditions of cellular stress, a Tet-inducible EGFP-RBM45 construct was generated by inserting the EGFP-RBM45 (N-terminal EGFP fusion) into the pcDNA4/TO vector (Invitrogen). The sequences of all constructs were confirmed by Sanger DNA sequencing and the sizes of the expressed proteins confirmed by Western blot.

### Inducible EGFP-RBM45 Cell Line

#### Creation of the Tet-On Parental Stable Cell Line

HEK293 cells were transfected with pcDNA6/TR (Invitrogen) to express the Tet repressor and selected with 10 μg/ml Blasticidin (Invivogen, San Diego, CA, USA). Multiple stable clones were isolated, propagated, transfected with pcDNA4/TO/lacZ (Invitrogen), and induced with 10 μg/ml doxycycline hyclate (Sigma) for the expression of ß-galactosidase. The ß-Gal assay was performed to screen for clones with the highest expression of the Tet repressor. A Tet-on parental clone with a combination of the lowest ß-galactosidase levels without doxycycline induction and highest expression of ß-galactosidase in the presence of doxycycline was then selected. This line was maintained with 5 μg/ml Blasticidin.

#### Creation of Inducible EGFP-RBM45 Cell Line

pcDNA4/TO-EGFP-RBM45 was transfected into the Tet-on parental stable cell line above and EGFP-positive stable cell lines were selected in the presence of 100μg/ml Zeocin.

### CRISPR-Cas9 Modified FLAG-RBM45 Cell Line

A 3xFLAG tag was appended to the N-terminus of the endogenous RBM45 genomic locus in HEK293 cells using the CRISPR-Cas9 method described by Ran et al^38^ with some modifications. <u>sgRNA preparation:</u> Five sgRNA sequences near the N-terminus of the RBM45 genomic locus were selected using the CRISPR Design Tool (http://crispr.mit.edu)^39^. Oligos encoding the five sgRNAs were individually cloned into the pSpCas9(BB)-2A-Puro (PX459) V2.0 vector (Addgene #62988) and transfected into HEK293 cells. Cells were selected in the presence of 5 µg/ml puromycin (Invivogen) for 48 hours before functional validation of sgRNAs. The cleavage efficiency of sgRNAs were evaluated by the TIDE (Tracking of Indels by DEcomposition) assay^40^. Briefly, genomic DNA from puromycin selected cells and wild-type HEK293 cells was extracted (Promega Wizard Genomic DNA Purification Kit #A1120; Promega, Madison, WI, USA) for PCR amplification of a 535 bp region flanking the sgRNA cleavage site. The PCR DNAs were gel purified (Promega Wizard SV Gel and PCR Clean-up System #A9282) and 30 ng of each PCR DNA was Sanger sequenced. The sequencing result for each sgRNA was aligned to the wild-type control by TIDE software (https://tide.nki.nl/) and the resulting traces were used to calculate cleavage efficiencies^40^. The two sgRNAs with the highest cleavage efficiency (both 85%) were chosen for co-transfection with a homology-directed repair (HDR) template for CRISPR-Cas9 genome editing.

#### Generation of CRISPR-Cas9 Modified FLAG-RBM45 HEK293 Cell Line

A 200nt ssODN ultramer (IDT) was used as a homology-directed repair (HDR) template for CRISPR-Cas9 editing of the endogenous RBM45 locus. Co-transfection of 500 ng sgRNA plasmid and 1 µl of 10 µM ssODN template into 2×10^5^ HEK293 cells was performed using nucleofection on the 4D-Nucleofector system (LONZA, Basel, Switzerland), as described previously^38^. The transfected cells were enriched by selection with 1µg/ml puromycin for 48 hours. Clonal cell lines were isolated by serial dilution and expanded for 3 weeks. Genomic DNA from cell lines growing on 96-well plates was extracted by 25 µl QuickExtract DNA extraction solution (Epicentre #QE09050, Madison, WI, USA), further diluted to 200 µl with H_2_O. CRISPR-Cas9 genome editing was assessed by using genomic DNA from these samples for PCR amplification of the RBM45 genomic locus using a FLAG-specific forward primer and an RBM45 reverse primer flanking a 480 bp region. The 480 bp PCR DNAs were then Sanger sequenced to verify the correct insertion of the 3xFLAG into the desired genomic locus. Immunostaining and real-time PCR were performed to assess the nuclear morphology, subcellular localization, and expression level of CRISPR modified FLAG-RBM45 protein and mRNA expression levels as compared to wild-type HEK293 cells.

#### Primers Used for TIDE Assay

PCR forward primer: 5’-GGACTCCTCTTTCTCCCGGAAGCGG

PCR reverse primer: 5’-GCTGCCGGGAACGGATGTTCCACTC

Sequencing forward primer: 5’-ACTCCTCTTTCTCCCGGAAGCGGAG

Sequencing reverse primer: 5’-TGCCGGGAACGGATGTTCCACTCCT

#### Target Sequences of the Two sgRNAs with the Highest Cleavage Efficiency

5’-TGCATTCGGGTGGAGCACCA (sense strand)

5’-AGAGCTGCCAGCTTCGTCCA (anti-sense strand)

#### Sequence of ssODN

Uppercase letters indicate the sequence for the 2xFLAG tag:

5’-

gagacagcagcggtggcagacaccgcagaagcaaagagcagtgaggctcctgcattcgggtggagcaccatgGACTACAAA GACGATGACGACAAGGACTACAAGGATGACGATGACAAAGCTGCTgacgaagctggcagctctgcgagcggcgggggcttccgcccgggcgtggacagcctggacgaaccgcccaaca

#### Primers to PCR Amplify CRISPR Modified Endogenous FLAG-RBM45

FLAG-specific forward primer: 5’-TCGGGTGGAGCACCATGGACTACAAAG

RBM45 reverse primer: 5’-GCTGCCGGGAACGGATGTTCCACTC

### siRNA knockdown of RBM45 and SatIII

To assess the effects of knockdown of endogenous RBM45 or SatIII expression on NSB formation, siRNAs targeting each transcript were designed based on previous studies^41, 42^ and transfected into HEK293 cells. Two siRNAs targeting RBM45 and one siRNA targeting SatIII were used, along with a scrambled siRNA as a negative control. For all siRNA knockdown studies, 40% confluent HEK293 cells growing on 24- or 96-well plates were transfected with 15 pmol or 3 pmol of siRNA, respectively, using the Lipofectamine 3000 reagent (Invitrogen) according to the manufacturer’s instructions. Cells were immunostained or assayed for cellular viability 48 hours post-transfection. siRNAs (Dharmacon, Layfayette, CO, USA) with the following target sequences were used:

RBM45-1: 5’-GUAUGGAGAUAUCGAGUAU

RBM45-2: 5’-GGACAUGAACCUAGAGUAA

SatIII: 5’-UGGAAUGGAAUGGAAUGGA

The negative control scrambled siRNA (Dharmacon # D-001810-10-05) consists of a pool of 4 sequences previously determined to have minimal effects on gene expression in mammalian cells^43^. Real-time PCR was used to quantify the siRNA knockdown efficiency for each transcript. cDNAs were made with SuperScript VILO Mastermix (Invitrogen) containing the random hexamer and OligodT primers. The relative quantity was normalized with GAPDH and calculated using the ΔΔCt method. The primers for real-time PCR were:

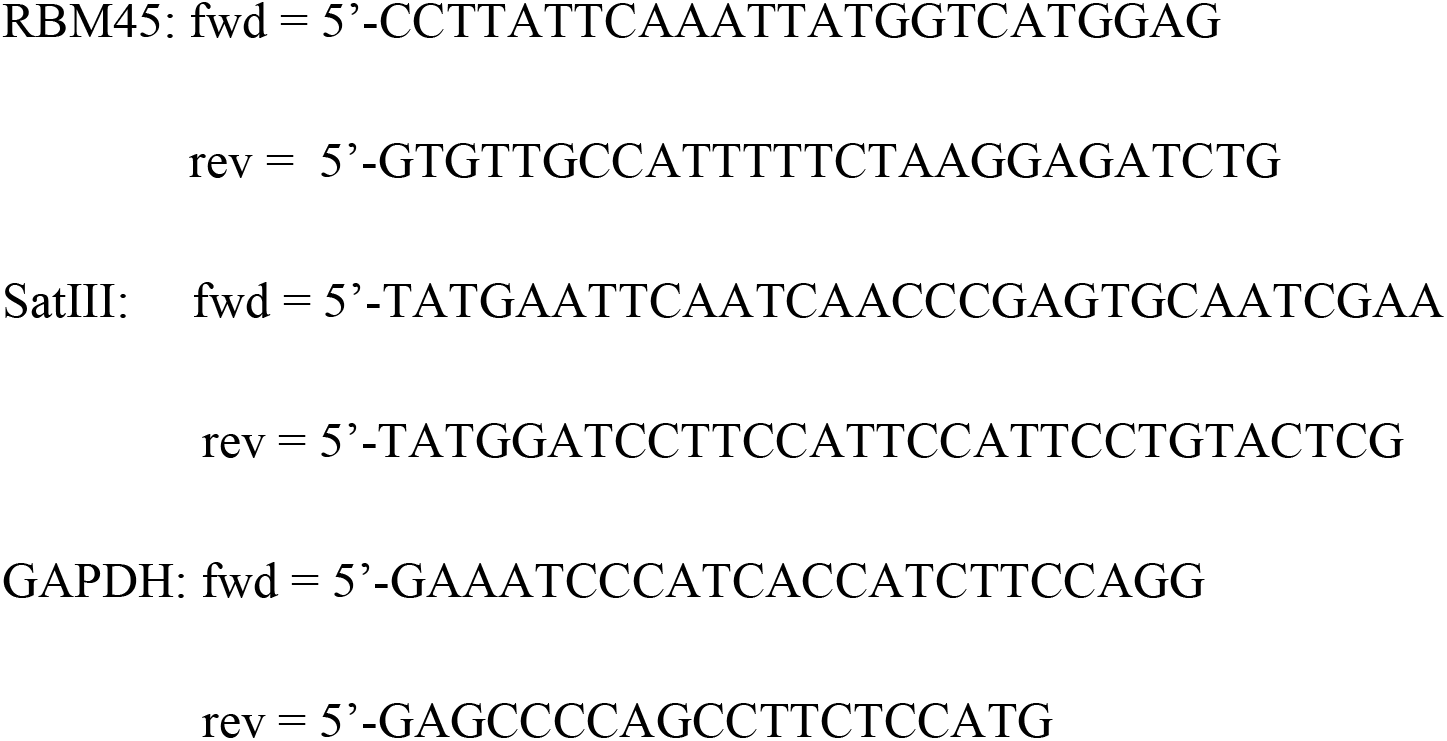

### Subcellular Fractionation and Assessment of RBM45 Solubility

HEK293 cells were grown in 10 cm tissue culture dishes for fractionation into nuclear/cytoplasmic or total protein extracts. All extraction procedures were done at 4°C in the presence of a protease inhibitor cocktail (Sigma). Cytoplasmic extracts were obtained by immersing cells in a hypotonic buffer (10 mM HEPES, 1.5 mM MgCl_2_, 10 mM KCl, 0.5 mM DTT) for 5 minutes and then lysing cells by homogenization with a Dounce homogenizer. The lysed cells were spun for 5 minutes at 1,000 rpm to pellet nuclei and the supernatant was retained as the cytoplasmic fraction and solubilized in RIPA buffer. Cells were visualized by brightfield microscopy to confirm hypotonic buffer-induced swelling, cell membrane lysis, and nuclear membrane integrity following homogenization. The pelleted nuclei were re-suspended, centrifuged at 2800 g for 10 minutes, and broken with sonication, which was also monitored by brightfield microscopy. Following centrifugation at 2800 g for 10 minutes, the supernatant was retained as the nuclear fraction and solubilized in RIPA buffer. The purity of the lysate was determined by SDS-PAGE and Western blotting for GAPDH and lamin A/C as markers of the cytoplasm and nucleus, respectively.

For assays examining the solubility of RBM45 under various stressor conditions, HEK293 cells constitutively expressing FLAG-RBM45 (overexpressed or CRISPR-modified, as described above) were grown on 10 cm plates until confluent and were treated as indicated to induce the formation of NSBs. The cells were harvested and fractionated into soluble and insoluble protein fractions as previously described^44^ with minor modifications. Cells from one 10 cm plate were harvested, lysed in 1.5ml RIPA buffer (150 mM NaCl, 1% NP40, 0.1% SDS, 50 mM Tris, pH 8.0) at 4°C for 15 min, sonicated, and centrifuged at 100,000g for 30 min at 4°C. After centrifugation, the supernatant was collected as the RIPA-soluble fraction and its protein concentration was determined by BCA assay (Pierce). To prevent carryover of residual soluble protein extract into the insoluble fraction, the resulting pellets were washed once with RIPA buffer, resonicated, recentrifuged at 100,000g for 30 min at 4°C, and the supernatant was removed. RIPA-insoluble pellets were then extracted in 1.5 ml urea buffer (7 M urea, 2 M thiourea, 4% CHAPS, 30 mM Tris, pH 8.5) at room temperature for 20 min, sonicated, and centrifuged at 100,000 g for 30 min at 22°C. The final supernatant was collected as the insoluble, detergent-resistant fraction and its protein concentration was determined by Bradford assay (Bio-Rad, Hercules, CA, USA). To inhibit protein degradation, 1 mM PMSF (a protease inhibitor) was added to all buffers prior to use.

### SDS-PAGE and Western Blotting

SDS-PAGE, total protein staining, and Western blotting were performed as previously described^45^. In brief, samples were mixed with LDS sample buffer (Invitrogen), DTT reducing reagent, and deionized water (as a diluent), heated at 70°C for 10 minutes, and loaded onto 1.5 mm 4-12% Bis-Tris gradient SDS-PAGE gels (Invitrogen). For assays examining HSF1, SAFB, and RBM45 abundance, 10 µg of protein was loaded from lysates prepared as above. For assays examining the solubility of RBM45, 15 μg of the RIPA-soluble fraction and 4 μg of the detergent-resistant fraction were loaded in each lane. Gel runs were performed in 1X MOPS buffer at a constant 200 V at 4°C. Following SDS-PAGE, proteins were transferred from gels to PVDF membranes. Wet tank transfer was performed in Towbin buffer (25 mM Tris, 192 mM glycine) using a ramped transfer approach, as in Otter et al.^46^, where samples were transferred for 6 hours at a constant 8 V, then the voltage increased to 16 V for a further 6 hours. Following the transfer, membranes were dried for 1 hour, then rehydrated using methanol and deionized water.

For Western blotting, membranes were initially blocked in Odyssey blocking buffer (Licor) for 60 minutes. Membranes were incubated in primary antibodies for 1-12 hours and washed three times for 15 minutes each in a phosphate buffered saline (PBS) solution with Odyssey blocking buffer added at a dilution of 1:10. Following the washing step, membranes were incubated in solutions containing secondary antibodies conjugated to IRDye 680 and IRDye 800 (LI-COR, Lincoln, NE, USA) for 60 minutes and washed as above. After secondary antibody incubations membranes were washed 3 times for 15 minutes each in Odyssey blocking buffer diluted 1:10 in PBS. Finally, immediately prior to imaging, membranes were washed once in PBS to remove residual detergent from the membrane. Blot images were acquired on an Odyssey CLx scanner (LI-COR) at 169 µm resolution at the highest intensity setting at which no pixel saturation was present in the resulting image.

### Immunocytochemistry and Image Analysis

Indirect immunofluorescence was performed as in Li et al.^5^ In brief, cells were grown on 20 mM #1.5 PDL-coated coverslips. When cells reached 70% confluence, they were treated as indicated, washed with 1X PBS, fixed in 4% paraformaldehyde for 10 minutes, washed 3 times in PBS, and permeabilized for 10 minutes using 0.1% Triton X-100 in PBS. Following additional washing with 1X PBS to remove fixative and detergent, cells were blocked with Superblock (Scytek, Logan, UT, USA) for 1 hour and immersed in primary antibody solutions for 2 hours. Subsequently, they were washed 4 times (10 minutes each) with IF wash buffer (1:10 Superblock:1X PBS), and immersed in secondary antibody solutions for 1 hour. Secondary antibodies used were goat-anti-rabbit Cy2 (Abcam, Cambridge, UK) and goat-anti-mouse Cy5 (Abcam). Following secondary antibody incubations, coverslips were washed 4 times as above, washed 4 times with 1X PBS, incubated in a 300 nM DAPI solution for 10 minutes, and washed 4 times with 1X PBS. Coverslips were immersed in increasing concentrations of 2,2’-thiodiethanol (TDE) according to the method of Staudt et al.^47^, and mounted in a 97% TDE solution with a refractive index of 1.518 to match that of the immersion oil used for imaging.

Slides were imaged on a Observer Z1 (Zeiss, Jena, Germany) microscope with a 1.4 NA 63x objective with LED illumination. All images were acquired as three-dimensional Z-stacks with a Z sampling depth of 12 μM, X/Y sampling interval of 0.102 μM, and a Z sampling interval of 0.240 μM. All images were subjected to shading correction and background subtraction prior to deconvolution. Following image acquisition and initial processing, images were deconvolved using Huygens digital deconvolution software (SVI). A measured PSF was generated by imaging 200 nm diameter fluorescent Tetraspeck beads (a sub-resolution particle in this imaging system) (Life Technologies, Waltham, MA, USA) mounted in 97% TDE and inputting the obtained images into the Huygens software’s PSF distiller application. The performance of the measured PSF was assessed by deconvolving additional 10 μm image stacks of 200 nm fluorescent beads using the measured and theoretical PSFs. Comparisons of the deconvolution result using the different PSFs demonstrated that the measured PSF provided a superior image restoration result as assessed by the size and shape of the deconvolved bead image. Alignment of individual channels in the measured PSF was then used to correct images for spherical aberration. All cell images were then deconvolved using the measured PSF and a maximum likelihood deconvolution algorithm.

### Immunohistochemistry and Image Analysis

Immunohistochemistry of lumbar spinal cord and hippocampal brain tissue sections from non-neurological disease control, sporadic ALS (sALS), c9ORF72 hexanucleotide repeat expansion ALS (c9ORF72), and non-c9ORF72 linked familial ALS (fALS), frontotemporal lobar degeneration (FTLD), and Alzheimer’s disease (AD) subjects was performed as previously described^1^. Tissue sections were obtained from the Barrow Neurological Institute and Target ALS Multicenter post mortem tissue bank cores. All participants in the post mortem tissue bank cores provided IRB approved informed consent for the collection of post mortem tissues. Subject demographics are shown in Table 1. Formalin fixed, paraffin embedded tissue sections (6 μm thick) were deparaffinized by immersion in xylene, rehydrated by successive immersion in increasing concentrations of ultrapure water in ethanol, and subjected to antigen retrieval. Antigen retrieval was performed in citra buffer (pH 6.0; Biogenex, Fremont, CA, USA) by steam heating for 30 minutes. Sections were washed in PBS, blocked using Superblock (Scytek), and immersed in primary antibody solutions overnight. The next day, the slides were washed 4 times in PBS and incubated in solutions containing appropriate secondary antibodies conjugated to Alexafluor 488 and 594 for 1 hour. Subsequently, slides were again washed 4 times in PBS, and immersed in a 300 nM DAPI solution for 10 minutes to stain cell nuclei. Following PBS washing, slides were immersed in a Sudan black-based autofluorescence eliminator reagent (Millipore) to quench endogenous lipofuscin autofluorescence and mounted in gelvatol.

**Table 1.** Subject Demographics. Demographics for each case are shown. PMI = post mortem interval (hours between death and tissue harvest), C9ORF72 = presence or absence of C9ORF72 hexanucleotide repeat expansion.

| Subject | Group | c9orf72 | Age | Sex | PMI | Hippocampus | Spinal Cord |
| --- | --- | --- | --- | --- | --- | --- | --- |
| 1 | Control | no | 54 | M | 6 | yes | yes |
| 2 | Control | no | 53 | F | 4 | yes | yes |
| 3 | Control | no | 51 | F | 5 | yes | yes |
| 4 | Control | no | 76 | M | 13 | yes | yes |
| 5 | Control | no | 57 | M | 2 | yes | yes |
| 6 | Control | no | 48 | M | 2 | yes | yes |
| 7 | Control | no | 58 | F | 5 | yes | yes |
| 8 | Control | no | 76 | M | 14 | yes | no |
| 9 | Control | no | 82 | F | 5 | yes | yes |
| 10 | Control | no | 57 | F | 11 | yes | yes |
| 11 | ALS | no | 62 | M | 7 | no | yes |
| 12 | ALS | no | 50 | F | 7 | no | yes |
| 13 | ALS | no | 60 | F | 4 | no | yes |
| 14 | ALS | no | 59 | M | 4 | yes | yes |
| 15 | ALS | no | 77 | F | 4 | yes | yes |
| 16 | ALS | no | 79 | F | 4 | yes | yes |
| 17 | ALS | no | 52 | F | 21 | yes | yes |
| 18 | ALS | no | 45 | M | 7 | yes | no |
| 19 | ALS | no | 65 | M | 5 | yes | no |
| 20 | ALS | no | 66 | M | 18 | yes | yes |
| 21 | ALS | no | 53 | F | 12 | yes | yes |
| 22 | ALS | no | 73 | F | 6 | no | yes |
| 23 | ALS | no | 59 | M | 5 | no | yes |
| 24 | ALS | yes | 68 | M | 4 | no | yes |
| 25 | ALS | yes | 43 | M | 6 | no | yes |
| 26 | ALS | yes | 63 | F | 4 | no | yes |
| 27 | ALS | no | 48 | F | 5 | no | yes |
| 28 | AD | no | 71 | M | 4 | yes | no |
| 29 | AD | no | 84 | F | 5 | yes | no |
| 30 | AD | no | 85 | M | 5 | yes | no |
| 31 | AD | no | 73 | F | 4 | yes | no |
| 32 | AD | no | 71 | F | 5 | yes | no |
| 33 | FTLD | no | 69 | M | 6 | yes | no |
| 34 | FTLD | no | 67 | M | >24 | yes | no |
| 35 | FTLD | no | 69 | M | 6 | yes | no |
| 36 | FTLD | no | 85 | M | 3 | yes | no |
| 37 | FTLD | no | 91 | F | 8 | yes | no |
| 38 | FTLD | no | 79 | M | 7 | yes | no |

All tissue immunofluorescence images were acquired on a Zeiss LSM 720 confocal microscope. Images were acquired as three-dimensional Z-stacks with a Z sampling range of 15 μm, X/Y sampling interval of 0.102 μM, and a Z sampling interval of 0.240 μm. All images were subjected to shading correction and background subtraction, and image processing was performed using NIH ImageJ^48^. We developed an image analysis pipeline to (1) count the number of cells in hippocampal dentate gyrus and lumbar spinal cord images, (2) count the number of RBM45 nuclear inclusions in each cell in these images, and (3) measure the nuclear immunoreactivity of SAFB in each cell. These steps were carried out as follows. To count the number of cells in each image, we created binary images of DAPI-stained nuclei using fixed intensity thresholding. The ImageJ particle analyzer was then used to count the number of DAPI-positive cells in each image. Cell counts were further filtered by excluding possible false-positives on the basis of cell size and shape. To count the number of RBM45 nuclear inclusions, we used the DAPI binary images as an overlay mask to restrict analysis to RBM45 foci in the nuclei of cells. These masked images were then thresholded using the Robust Automatic Threshold plugin^49^. The ImageJ particle analyzer was then used to count the number of RBM45-positive foci in the nucleus with particle sizes restricted to published parameters for the size of nuclear stress bodies^33, 50^. Finally, the SAFB immunoreactivity of each nucleus was measured using the corresponding DAPI binary image as an overlay mask. Each step in the image analysis pipeline was evaluated by comparing its performance to manual counts of at least 10 independent images, with a requirement of 95% agreement between automated and manual counts across images. For each subject, a total of 8 fields per region (hippocampal dentate gyrus or lumbar spinal cord) were analyzed.

### Antibodies

The following primary antibodies and dilutions were used for all Western blotting experiments, rabbit polyclonal anti-C-terminal RBM45 (Pacific Immunology, Fremont, CA, USA, custom, 1:2,000), rabbit polyclonal anti-N-terminal RBM45 (Pacific Immunology, custom, 1:2,000), rabbit polyclonal anti-RBM45 (Sigma, 1:1,000), mouse-monoclonal anti-HA (Sigma, 1:5,000), mouse monoclonal anti-FLAG M2 (Sigma F3165, 1:5,000), rabbit polyclonal anti-TDP-43 (Proteintech, Rosemont, IL, USA, 1:3000), mouse monoclonal anti-actin (Abcam, 1:5,000), rabbit monoclonal anti-lamin A/C (Epitomics, San Francisco, CA, USA, 1:1000), and rabbit monoclonal anti-GAPDH (Cell Signaling, Danvers, MA, USA, 2118S, 1:5,000). The following secondary antibodies were used for Western blotting experiments (each at a dilution of 1:10,000): goat anti-rabbit IRDye 680, goat anti-rabbit IRDye 800, goat anti-mouse IRDye 680, and goat anti-mouse IRDye 800 (LI-COR).

The following primary antibodies and dilutions were used for immunofluorescence and immunohistochemistry: rabbit polyclonal anti-RBM45 (Sigma, 1:75), rabbit polyclonal anti-RBM45 (custom; Pacific Immunology, 1:100), mouse monoclonal anti-scaffold attachment factor B (SAFB, Lifespan Biosciences, Seattle, WA, USA, 1:100), mouse monoclonal anti-SAFB (Proteintech, 1:100), rabbit polyclonal anti-SAFB (Proteintech, 1:100), mouse-anti-HSF1 (Abcam, 1:100), rabbit polyclonal anti-heat shock factor 1 (HSF1; Proteintech, 1:100), mouse monoclonal anti-SC35 (Abcam, 1:300), mouse monoclonal anti-coilin (Abcam, 1:200), mouse monoclonal anti-SMN (Sigma, 1:300), mouse monoclonal anti-G3BP (Genentech, 1:100), mouse monoclonal anti-TDP-43 (Proteintech, 1:100), mouse monoclonal anti-FUS (Proteintech, 1:100), and mouse monoclonal anti-HA (Sigma, 1:1,000). Goat-anti-rabbit Cy2 and goat-anti-mouse Cy5 (Abcam) secondary antibodies were used for immunofluorescence at a dilution of 1:200. Goat-anti-rabbit Alexafluor 488 and goat-anti-mouse Alexafluor 594 (Life Technologies) secondary antibodies were used at a dilution of 1:200 for human tissue immunohistochemistry experiments.

### Data Analysis

The statistical analysis of all data was performed using Microsoft Excel (Microsoft, Redmond, WA, USA) and R. Between groups comparisons were performed using one-way analysis of variance (ANOVA) with Tukey’s HSD post-hoc test. For the analysis of RBM45 nuclear inclusions per cell in human tissues, we used the non-parametric Mann-Whitney U test to perform all possible two group comparisons. The Bonferroni correction was used to account for multiple testing in these comparisons. Linear regression was used to analyze the relationship between RBM45 nuclear inclusion number and SAFB nuclear immunoreactivity. All figures were constructed in Adobe Illustrator (Adobe, Mountain View, CA, USA) and Inkscape.

## Results

### RBM45 Associates with Nuclear Stress Bodies (NSBs)

To further define the subcellular distribution of RBM45 and the mechanisms leading to RBM45 nuclear inclusion formation, we assessed RBM45’s co-localization with markers of several membraneless organelles in HEK293 cells. We used established marker proteins for nuclear speckles (SC35), Cajal bodies (coilin), nuclear gems (SMN), stress granules (SGs; G3BP), and nuclear stress bodies (NSBs; SAFB and HSF1). As shown in Figure 1A-C, under basal conditions, RBM45 is a predominantly nuclear protein with minimal cytoplasmic immunoreactivity. Moreover, the protein is diffusely localized throughout the nucleus and does not co-localize with markers for nuclear speckles, Cajal bodies, or nuclear gems.

**Figure 1.**
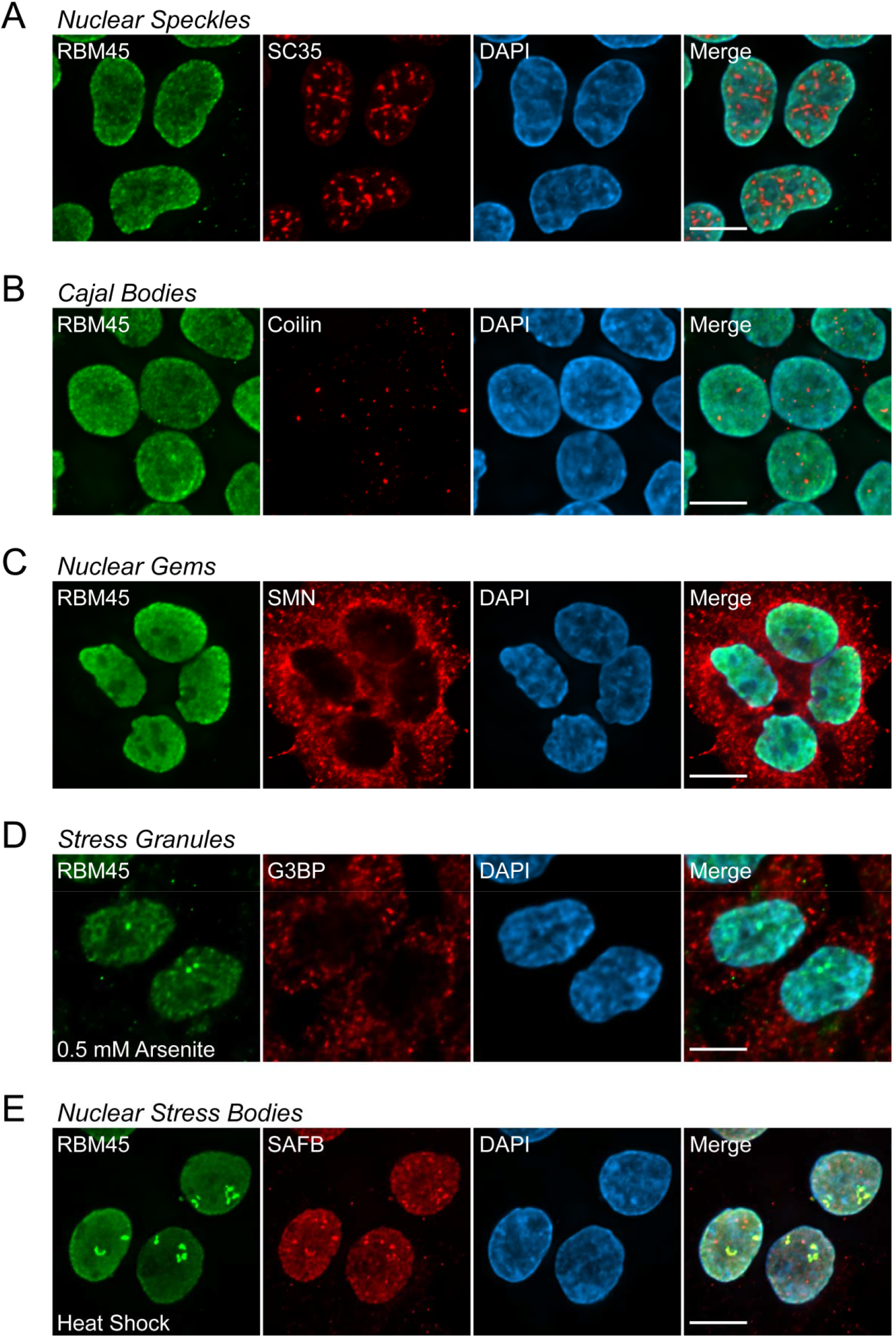
RBM45 Association with Nuclear Organelles. HEK293 cells were immunostained for endogenous RBM45 and marker proteins of the indicated nuclear structures. Under basal conditions, RBM45 is diffusely localized throughout the nucleus and does not associate with nuclear speckles, Cajal bodies, or nuclear gems (A-C, respectively). Following the onset of cellular stress (0.5 mM sodium arsenite, 1 hour [D] or heat shock at 42°C for 1 hour [E]), RBM45 redistributes to nuclear foci. (D) RBM45 does not associate with G3BP-positive cytoplasmic stress granules during cellular stress. (E) Stress-induced RBM45 nuclear foci correspond to NSBs, indicated by co-localization of RBM45 with the NSB marker protein SAFB. For all images, scale bar = 5 μm.

We next examined whether cellular stress promotes redistribution of RBM45 into membraneless organelles. Using sodium arsenite (0.5 mM for 1 hour) or heat shock (42°C for 1 hour) to induce cellular stress, we observed that RBM45 forms nuclear foci (Figure 1D and E, respectively). RBM45 stress-induced nuclear foci exhibit robust co-localization with SAFB-positive NSBs following heat shock, leading to reduced RBM45 immunoreactivity in areas that do not contain foci (Figure 2E). Treatment with sodium arsenite also led to the formation of numerous G3BP-positive cytoplasmic SGs, but these did not contain RBM45 (Figure 2D).

**Figure 2.**
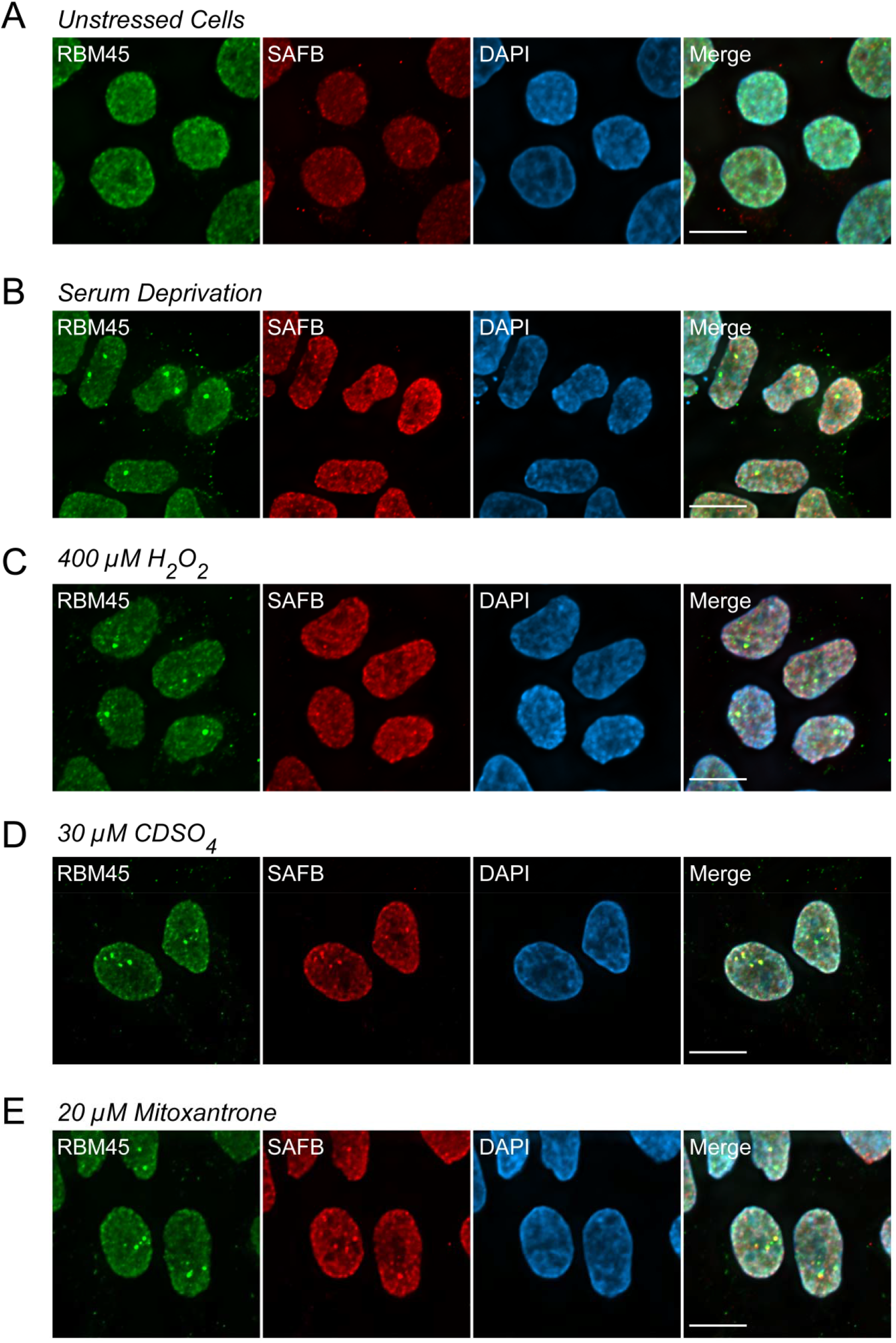
RBM45 Association with Nuclear Stress Bodies is a General Response to Cellular Stress. HEK293 cells were treated as indicated and the distribution of RBM45 and the NSB marker SAFB were evaluated by immunocytochemistry. (A) In untreated cells, the distribution of RBM45 and SAFB is diffuse and nuclear. (B-E) Treatment with the indicated stressors results in the formation of RBM45- and SAFB-positive NSBs. For all images, scale bar = 5 μm.

We then asked whether RBM45 association with NSBs occurs in response to other cellular stressors. In addition to sodium arsenite and heat shock (Figure 1E), we found that RBM45 coalesces into puncta and associates with NSBs under conditions of nutrient starvation (serum deprivation, Figure 2B), oxidative stress (400 µM H_2_O_2_, Figure 2C), heavy metal stress (30 µM CdSO_4_, Figure 2D), and genotoxic stress (20 µM mitoxantrone, Figure 2E). These results suggest that RBM45 association with NSBs is part of a general cellular stress response. As in our initial experiments, under basal conditions, the distribution of RBM45 was diffuse and nuclear, with no puncta or NSBs visible (Figure 2A).

NSBs are protein-RNA complexes that result from stress-induced transcription of pericentromeric heterochromatic satellite III (SatIII) repeats. SatIII transcripts remain in close proximity to their genomic locus and act as scaffolds for NSB assembly by recruiting various NSB RNA binding proteins, including heat shock factor (HSF1) and scaffold attachment factor B (SAFB). NSBs are hypothesized to result from a redistribution of existing nuclear RBPs, rather than an increase in RBP expression/protein levels. To assess whether cellular stress alters RBM45 protein levels, we performed Western blotting for RBM45 in HEK293 total protein extracts from unstressed HEK293 cells and cells subjected to oxidative and genotoxic stress. Neither stressor increased RBM45, HSF1, or SAFB protein levels in HEK293 cells (*p* > 0.05; Figure S1), consistent with existing models of NSB formation^32, 50^.

We also determined whether RBM45-interacting proteins are recruited to NSBs by virtue of their interactions with RBM45. TDP-43 and FUS physically interact with RBM45, are predominantly nuclear proteins under basal conditions, and have defined roles in the cellular response to stress. We therefore assessed whether TDP-43 or FUS can be incorporated into NSBs during cellular stress. Heat shock did not lead to the incorporation of either TDP-43 or FUS into RBM45-positive NSBs (Figure S2A). We overexpressed HA-tagged versions of full-length TDP-43 and FUS and evaluated their effects on NSB formation and their association with NSBs. As in the preceding experiments, we did not observe association of TDP-43 or FUS with NSBs (Figure S2B). We observed that overexpression of TDP-43, but not FUS, was sufficient to promote NSB formation, consistent with previous studies showing that altered TDP-43 expression leads to cellular stress (Figure S2B)^51, 52^.

We then tested whether RBM45 is required for the formation of NSBs using siRNA-based knockdown of RBM45. qPCR experiments established that our RBM45 siRNAs effectively targeted RBM45, resulting in an approximately 70% decrease in transcript abundance (Figure S3). As a positive control, we evaluated the effects of SatIII transcript knockdown by siRNA on NSB formation, as prior studies have established that SatIII knockdown abrogates NSB formation^41, 42^. siRNA targeting of SatIII reduced transcript levels approximately 80% (Figure S3). Neither the scrambled siRNA nor knockdown of RBM45 prevented the formation of NSBs (Figure S4A, B, respectively). Consistent with previous findings, knockdown of SatIII transcripts prevented NSB formation and we found that the absence of NSBs was sufficient to prevent redistribution of RBM45, SAFB, and HSF1 to nuclear foci during cellular stress (Figure S4C). Therefore, RBM45 associates with NSBs during conditions of cellular stress, but is not essential for NSB formation.

**Figure 3.**
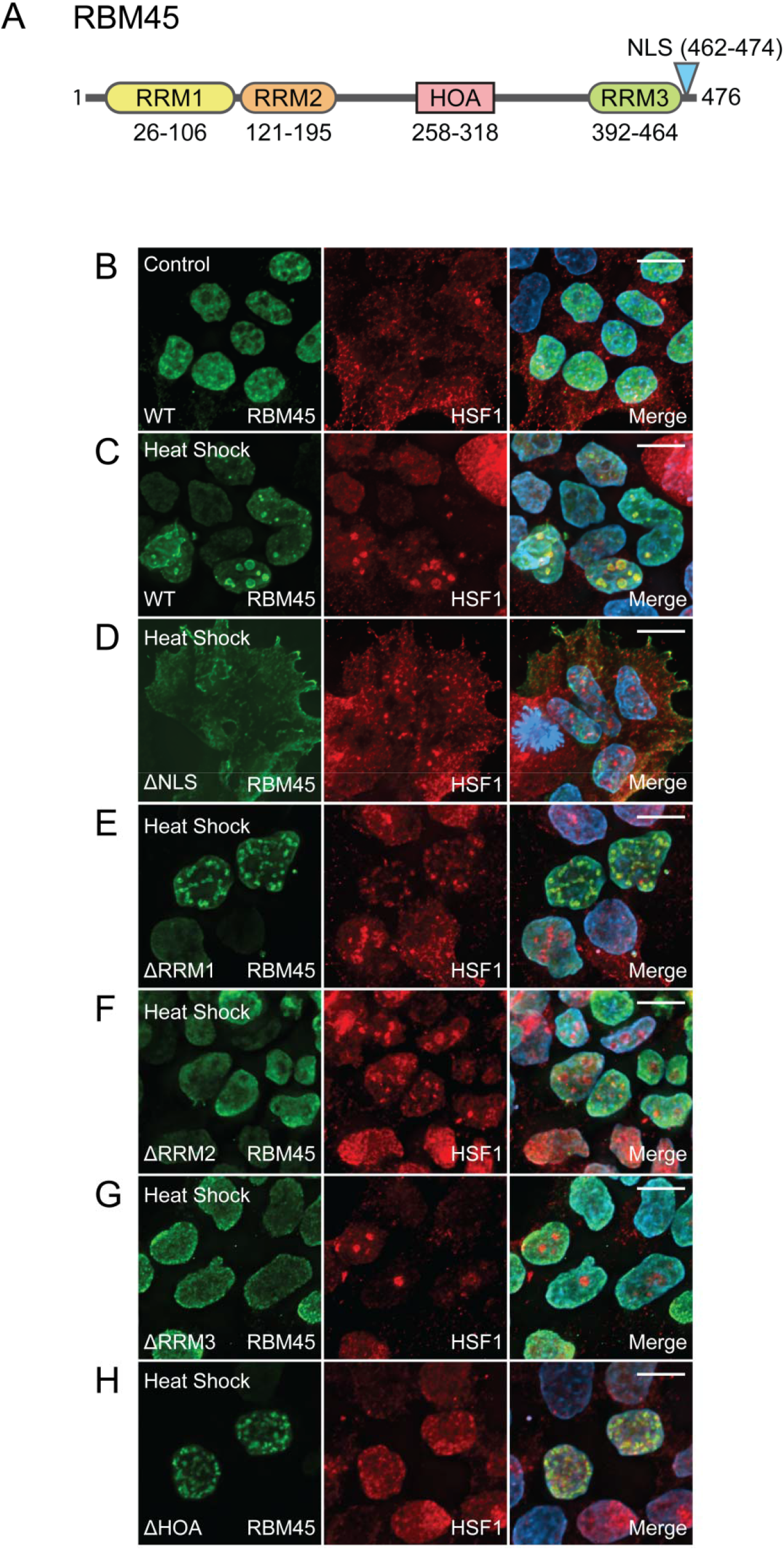
Mapping the RBM45 Domains Required for NSB Incorporation. (A) Schematic showing functional domains and their position in the full-length RBM45 protein. RRM = RNA recognition motif, HOA = homo-oligomerization domain, NLS = nuclear localization sequence. HEK293 cells were transfected with constructs encoding HA-tagged wild-type (WT) or domain-modified forms of RBM45 as indicated to determine which domains of the protein are necessary for incorporation into NSBs. (B) In unstressed cells, the distribution of WT RBM45 is predominately diffuse and nuclear. (C) Heat shock (42°C for 2 hours) leads to the robust formation of NSBs that are positive for RBM45 and the NSB marker HSF1. (D) Removal of the RBM45 NLS leads to its sequestration in the cytoplasm and prevents its association with NSBs during heat shock. (E) Removal of RRM1 does not alter the association of RBM45 with NSBs. (F) Removal of RRM2 prevents the association of RBM45 with NSBs. (G) Removal of RRM3 prevents the association of RBM45 with NSBs. (H) Removal of the RBM45 homo-oligomerization domain (HOA) does not alter the association of RBM45 with NSBs. For all images, scale bar = 5 μm.

### RNA Binding Domains are Essential for RBM45 Association with NSBs

RBM45 is a 474 amino acid protein with 3 RNA recognition motifs (RRMs), a bipartite nuclear localization sequence (NLS), and a homo-oligomer assembly (HOA) domain that mediates its self-association and interaction with other proteins (Figure 3A)^3, 6^. To identify which RBM45 domains are required for association with NSBs, we examined the co-localization of NSBs and HA-tagged full-length RBM45, several domain deletion mutant RBM45 constructs, and a nuclear localization sequence (NLS)-disrupting RBM45 mutant construct following heat shock. In unstressed cells, full-length HA-tagged RBM45 exhibited a diffuse nuclear localization and no HSF1-positive NSBs were observed (Figure 3B). A similar result was observed under basal conditions for all domain deletion constructs tested except the ΔNLS construct (not shown). Heat shock led to the incorporation of full-length, wild-type RBM45 in HSF1-positive NSBs (Figure 3C). Disruption of the C-terminal bipartite NLS of RBM45 led to cytoplasmic sequestration of the protein and this was sufficient to abrogate the association of RBM45 with NSBs (Figure 3D). Removal of the N-terminal RRM (RRM1) did not alter the association of RBM45 and NSBs (Figure 3E), nor did removal of the protein’s HOA domain (Figure 3H). In contrast, removal of either RRM2 or RRM3 was sufficient to prevent RBM45 from associating with NSBs (Figure 3F and 3G). Our data thus suggest that RBM45 association with NSBs occurs via direct RNA binding, rather than by indirect association with other NSB proteins.

### Chronic Stress Promotes RBM45 Aggregation via Association with NSBs

We next asked whether the association with NSBs is sufficient to promote nuclear RBM45 inclusion formation. Nuclear RBM45 inclusions occur in ALS, FTLD, and AD^1, 2^ and cellular stress has previously been shown to cause the formation of insoluble inclusions containing other ALS/FTLD-linked RBPs such as TDP-43 and FUS^20, 23, 24^. The formation of insoluble RBP inclusions may result from the persistent association of these proteins with cytoplasmic SGs. An analogous process occurring via RBP association with NSBs during cellular stress could lead to the formation of RBM45 nuclear inclusions. To understand the effects of acute and chronic stress on the distribution and aggregation of RBM45, we used a variety of cellular stressors at differing concentrations and for varying durations. Following acute and chronic stressor treatments, we performed immunocytochemistry for endogenous RBM45 and NSBs. As in prior experiments, under basal conditions, RBM45 was diffusely localized throughout the nucleus and no NSBs were visible (Figure 4A, B). Acute treatment with the genotoxic stressor MTX (5 μM for 6 hours) led to the formation of RBM45-positive NSBs (Figure 4C). To chronically stress cells with MTX, we tested a range of MTX concentrations (0.5-5 μM), seeking to find the maximum dose that did not adversely affect cellular viability. Using a dose of 1 μM MTX for 24 hours, we found that chronic stress led to an increase in the size, staining intensity, and number of RBM45 foci in the nucleus of HEK293 cells (Figure 4D). These foci were no longer positive for NSB marker proteins. We used a similar approach to compare the effects of acute and chronic stress resulting from treatment with the cadmium sulfate (acute – 30 μM, 2 hours; chronic – 5 μM, 24 hours), and sodium arsenite (acute – 1 mM, 1 hour; chronic – 0.1 mM, 24 hours). With all stressors, we found that chronic stress led to the formation of large, persistent RBM45-positive nuclear inclusions that were negative for NSB marker proteins (Figure 4E-H).

**Figure 4.**
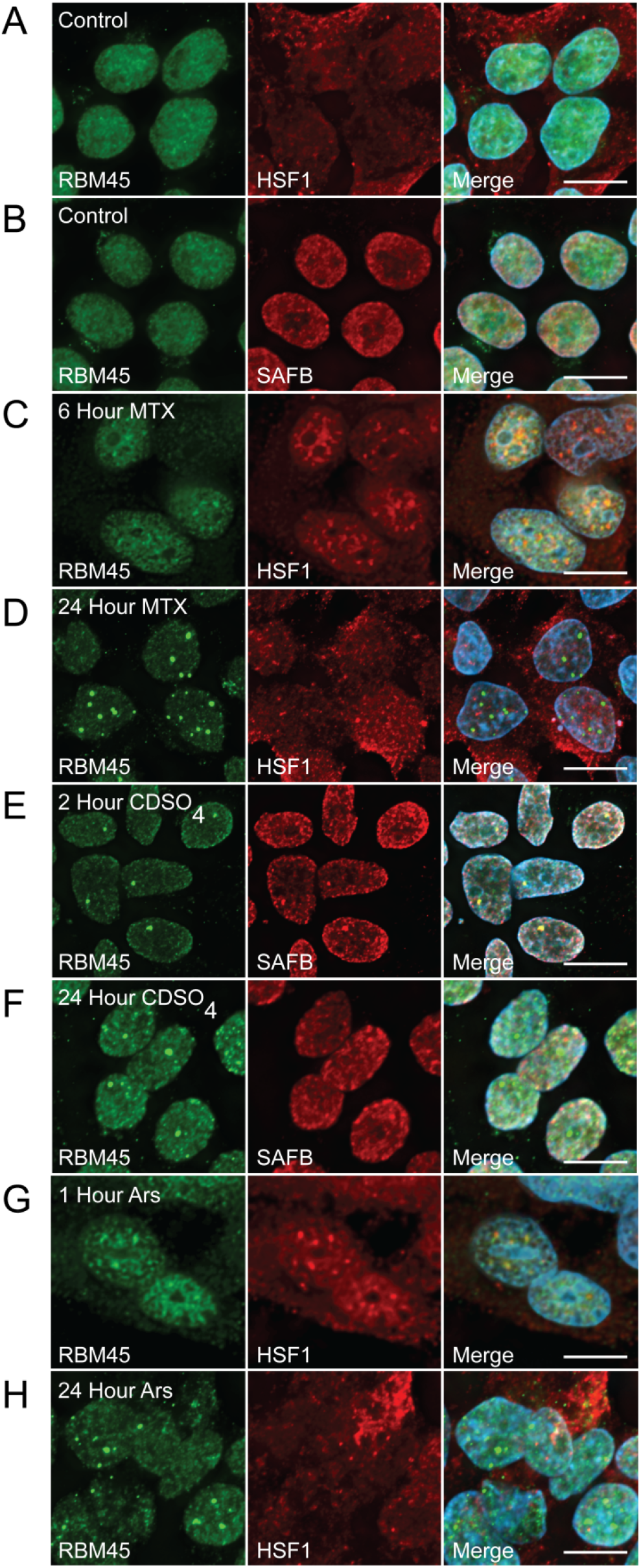
Chronic Stress Promotes RBM45 Nuclear Inclusion Formation. HEK293 cells were treated with three separate cellular stressors for varying durations to examine the effects of chronic and acute cellular stress on the subcellular distribution of RBM45 and nuclear stress body (NSB) formation. (A and B) In untreated cells, the distribution of RBM45 is diffuse and nuclear and no HSF1 (A) or SAFB (B)-positive NSBs are visible. (C) Acute treatment (6 hours) with the genotoxic stressor mitoxantrone (MTX; 5 µM) induces the formation of RBM45- and HSF1-positive NSBs. (D) Chronic treatment with MTX (24 hours; 1 µM) leads to the formation of RBM45 inclusions via NSB formation. At 24 hours, RBM45 nuclear inclusions are no longer positive for the NSB marker HSF1. (E and F) Similar results were obtained with acute (E) and chronic (F) treatment of cells with the oxidative stressor cadmium sulfate (CdSO4; 30 µM [acute] or 5 µM [chronic]) and the NSB marker SAFB. (G and H) Similar results were also obtained for acute and chronic treatment of HEK293 cells with sodium arsenite (Ars; 1 mM [acute; G] or 0.1 mM [chronic; H]). For all images, scale bar = 5 μm.

The RBM45 inclusions we observed following chronic stress may reflect a change in the solubility of RBM45. Because overexpression of proteins via transient transfection can lead to the formation of protein inclusions independent of cellular stress, we examined the solubility of endogenous RBM45 at physiological expression levels. For these experiments, we used CRISPR-Cas9 genome editing to append a 3x FLAG tag to the N-terminus of the endogenous RBM45 gene. We performed extensive characterization of CRISPR-Cas9-edited RBM45 HEK293 cells lines and found a non-significant reduction in RBM45 expression at the RNA level via qPCR in our CRISPR-RBM45 lines compared to unedited control cell lines (Figure S6A). We also observed no effects on nuclear morphology (Figure S6B), no differences in RBM45 subcellular localization (Figure S6B), and no effects on growth or cellular morphology (data not shown), suggesting that our CRISPR-cas9 genome editing and expression of HA-tagged RBM45 did not interfere with cellular physiology or RBM45 function. Soluble and insoluble protein extracts were obtained from FLAG-RBM45 cell lines as outlined in Figure 5A. The purity of the insoluble extract was determined by blotting for GAPDH, which is not a component of the insoluble protein fraction in stressed or unstressed cells. We used TDP-43, which increases in the insoluble fraction following cellular stress, as a positive control for our insoluble fraction Western blotting^44^. As shown in Figure 5B, heat shock or treatment with sodium arsenite led to a significant increase (*p* < 0.01 for both comparisons) in the amount of insoluble RBM45 (denoted using anti-FLAG tag antibody). For treatment with sodium arsenite, we observed that increasing the stress duration led to increased levels of insoluble RBM45 (*p* < 0.05). Treatment with sodium arsenite led to a concomitant decrease in the amount of soluble RBM45 protein (*p* < 0.01), while heat shock did not alter the levels of soluble RBM45 (*p* > 0.05). A similar pattern was observed for TDP-43, where sodium arsenite stress significantly increased the amount of insoluble TDP-43 and significantly decreased the amount of soluble TDP-43 compared to unstressed cells (*p* < 0.01 for both comparisons), while heat shock increased the amount of insoluble TDP-43 (*p <* 0.01), but did not alter the amount of soluble TDP-43 (*p* > 0.05).

**Figure 5.**
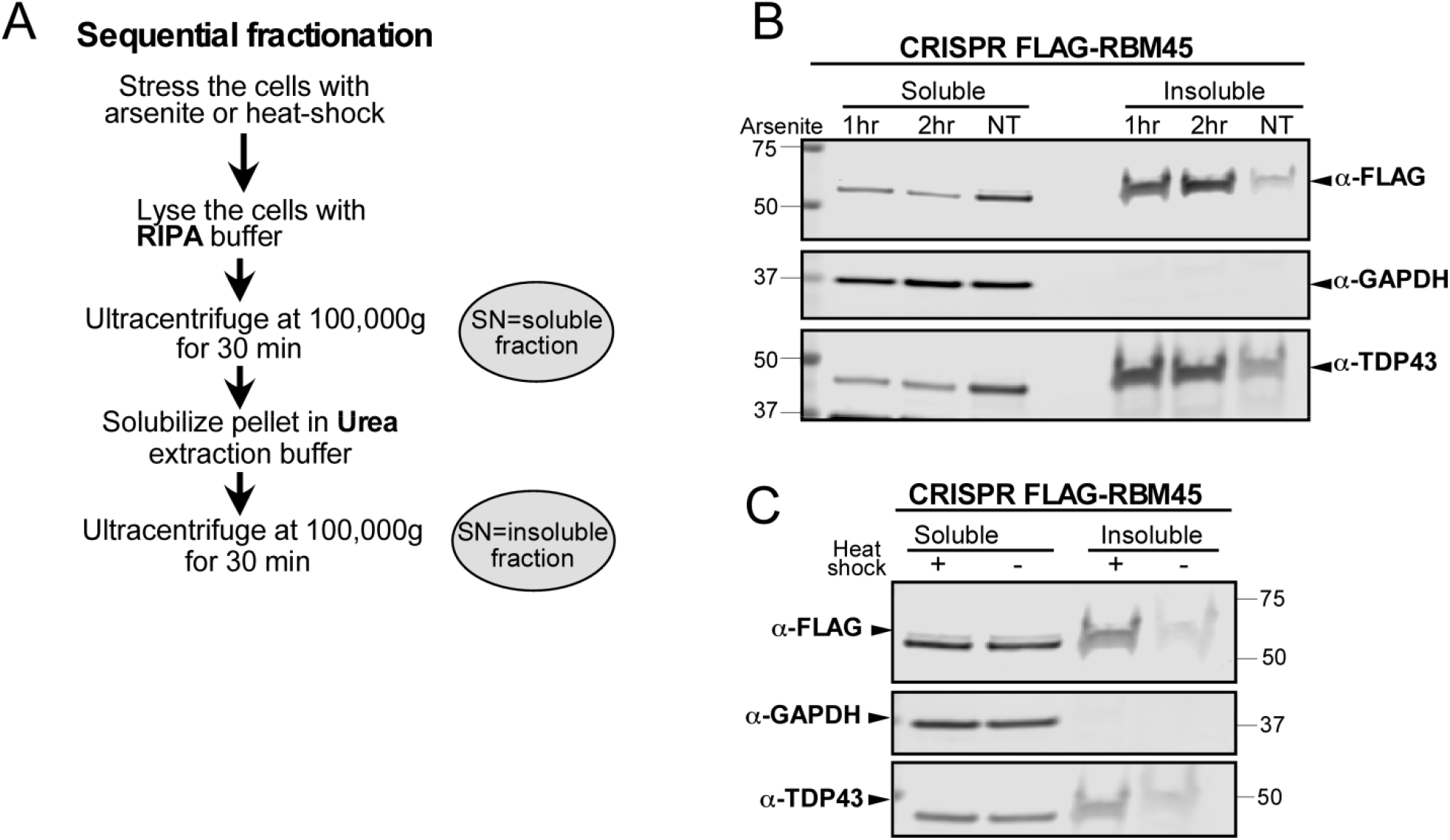
RBM45 Solubility during Conditions of Cellular Stress. (A) Workflow to obtain soluble and insoluble total protein extracts from stressed and unstressed HEK293 cells expressing FLAG-tagged RBM45. “SN” = supernatant. (B) Western blotting for FLAG-tagged RBM45 (α-FLAG), GAPDH (negative insoluble fraction control), and TDP-43 (positive insoluble fraction control) following sodium arsenite treatment (1 mM for 1 or 2 hours). (C) FLAG-RBM45, GAPDH and TDP-43 solubility following heat shock (1 hour at 42°C).

To understand the temporal dynamics of RBM45 NSB association and inclusion formation, we used live-cell imaging of HEK293 cells expressing N-terminally GFP-tagged RBM45 under the control of the Tet repressor. Addition of 10 μg/ml doxycycline to the culture medium led to robust expression of EGFP-RBM45. We used live-cell imaging to measure the dynamics of RBM45 during cellular stress. In untreated control cells, RBM45 maintained its diffuse nuclear localization for the 24-hour duration of the live-cell imaging experiments (Figure 6A). No RBM45 nuclear inclusions were observed during this period. In contrast, acute treatment with 0.5 mM sodium arsenite led to the formation of RBM45-positive NSBs (Figure 6B). These foci emerged within 1 hour of treatment (arrowheads in Figure 6B) and were almost completely eliminated 5 hours after removal of the stressor (Figure 6B, T=6 hr panel). At the completion of the experiment, no RBM45-positive NSBs were visible (Figure 6B, T=24 hr panel). In contrast, chronic treatment with a lower dose of sodium arsenite (0.1 mM for 12 hours) led to the formation of large nuclear RBM45 inclusions within 12 hours that persisted for the 24-hour duration of the experiment (Figure 6C, T=12 hr and T=24 hr panels). These inclusions were larger, more numerous, and brighter than those observed during acute stress conditions (compare Figure 6C T=12 hr panel to Figure 6B T=3 hr panel). The size and number of the inclusions increased over the course of the experiments and RBM45 NSBs did not disassociate, unlike in the acute stress condition (Figure 6C T=6, 12, and 24 hr panels). The presence of multiple, large RBM45 inclusions also impacted nuclear morphology (Figure 6C panel T=12 and T=24 hrs).

**Figure 6.**
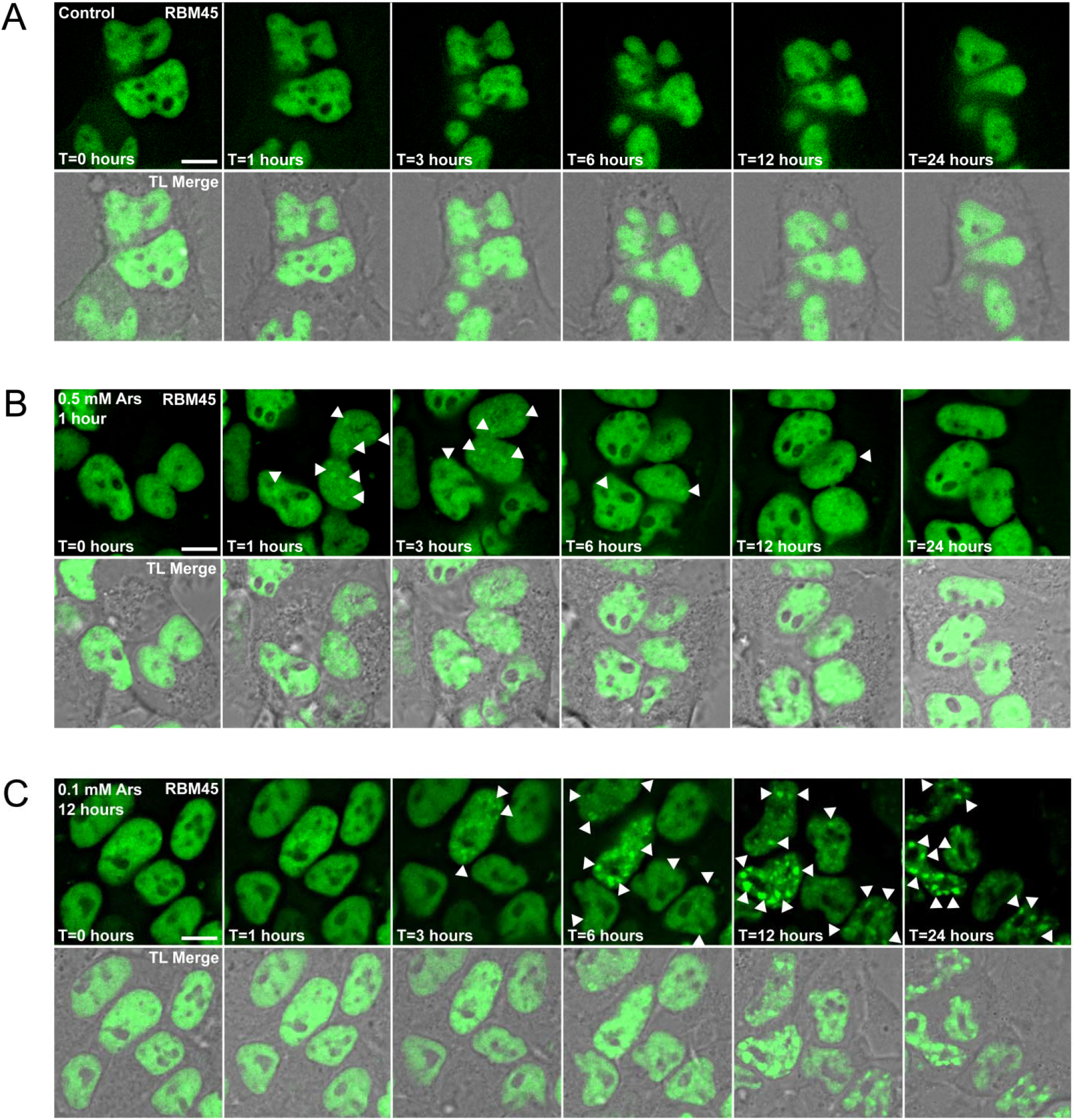
Live Cell Imaging of RBM45 during Conditions of Cellular Stress. HEK293 cells induced to express N-terminal GFP-tagged, full-length RBM45 were used to visualize the subcellular distribution of RBM45 during normal conditions and during cellular stress. (A) In the absence of stress, RBM45 is predominantly nuclear and diffusely localized throughout the nucleus. (B) Treatment with 0.5 mM sodium arsenite for 1 hour causes the formation of numerous RBM45-positive nuclear stress bodies (NSBs; arrowheads). Cells were treated with 0.5 mM sodium arsenite at t=0 hours. Under conditions of acute cellular stress, NSB formation is reversible, most RBM45-positive NSBs have dissipated by t=6 hours, and RBM45 remains diffusely localized throughout the nucleus for the remainder of the experiment. (C) Chronic cellular stress leads to the entrapment of RBM45 in nuclear inclusions. Cells were treated with 0.1 mM sodium arsenite for 12 hours to assess the effect of chronic cellular stress on RBM45 subcellular distribution. Because of the lower concentration of sodium arsenite, the time course of NSB formation is altered. Following removal of sodium arsenite, RBM45 remains in nuclear inclusions for the duration of the experiment. For all panels, scale bar = 5 μm.

### Quantification of RBM45 NSB Pathology in ALS, FTLD, and AD Subjects

We previously identified RBM45 nuclear and cytoplasmic inclusions in neurons and glia in ALS, FTLD, and AD^1^. To further characterize RBM45 pathology in these diseases, we sought to determine (1) to what extent nuclear RBM45 pathology resembles RBM45 inclusions resulting from chronic stress *in vitro*, (2) quantify RBM45 nuclear and cytoplasmic pathology across diseases, and (3) determine cell type patterns of RBM45 pathology. We first performed immunohistochemistry for RBM45, SAFB, and TDP-43 in the dentate gyrus of non-neurological disease controls (Figure 7A), FTLD (Figure 7B), ALS (Figure 7C), and AD (Figure 7D) subjects.

**Figure 7.**
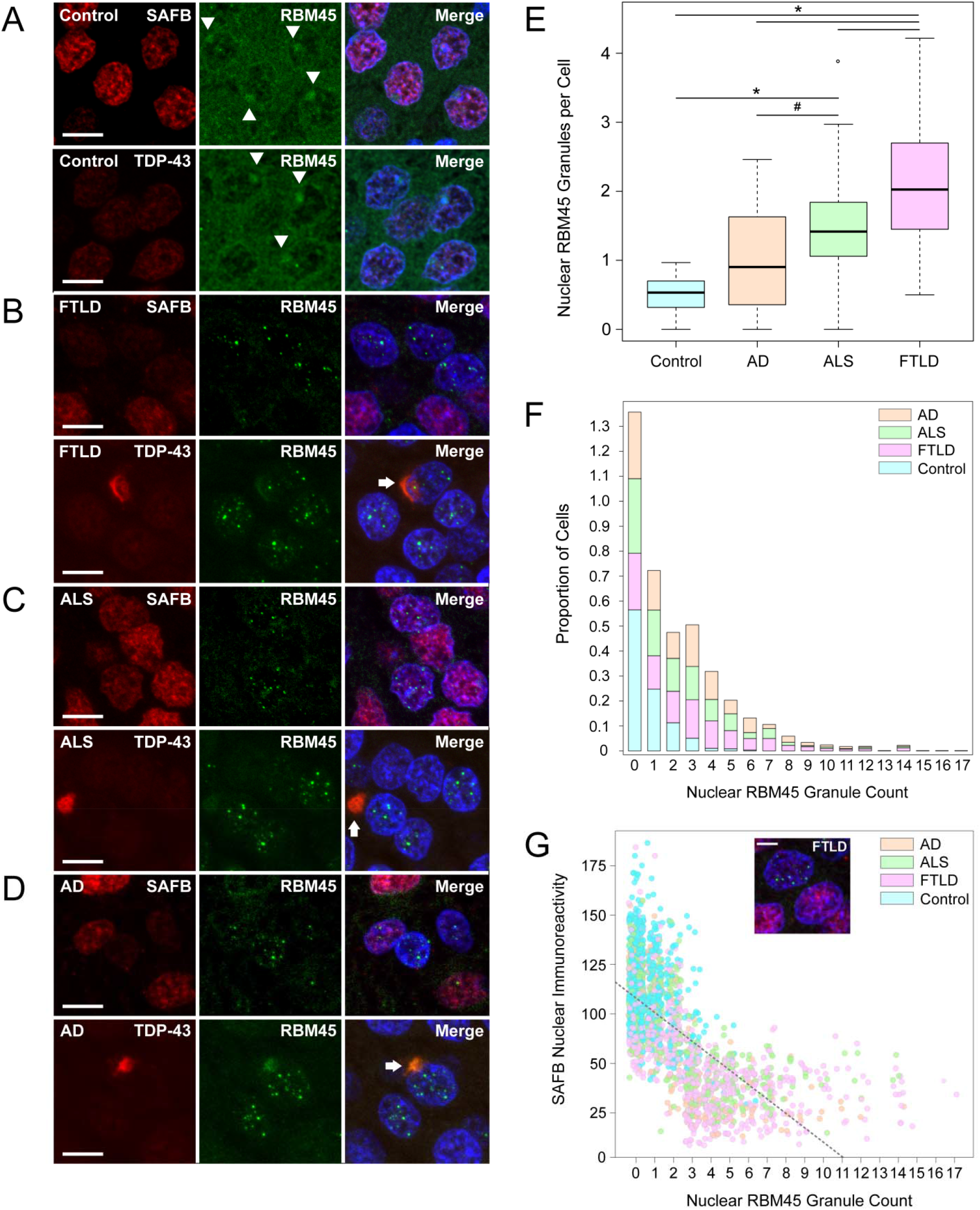
RBM45 Subcellular Distribution and Inclusion Pathology in FTLD, ALS, and AD. Immunohistochemistry for RBM45, SAFB, and TDP-43 was performed in hippocampal dentate gyrus from non-neurologic disease control, FTLD, ALS, and AD subjects. (A) Control subjects showed strong immunoreactivity for SAFB, TDP-43, and RBM45, with RBM45 immunoreactivity concentrated in the perinucleolar region (arrowheads) and SAFB and TDP-43 diffusely localized throughout the nucleus. (B) RBM45 immunoreactivity was predominantly in nuclear inclusions in FTLD dentate gryus cells, with attendant loss of cytoplasmic and perinucleolar immunoreactivity. Nuclear RBM45 inclusions lack SAFB and TDP-43. Dentate gyrus cells containing RBM45 nuclear inclusions also exhibit reduced SAFB immunoreactivity. Several TDP-43 and RBM45-positive cytoplasmic inclusions were found in FTLD subjects (arrow). (C) As in B, but for ALS subjects. Nuclear RBM45 inclusions lack SAFB or TDP-43, and cytoplasmic inclusions contain RBM45 and TDP-43 (arrow). (D) As in B, but for AD subjects. Nuclear RBM45 inclusions lack SAFB and TDP-43, and cytoplasmic inclusions contain RBM45 and TDP-43 (arrow). (E) Boxplot showing the number of RBM45 nuclear inclusions by group (# = *p* < 0.01, * = *p* < 1 × 10^-6^). (F) Stacked bar plot showing the number of RBM45 nuclear inclusions per cell by subject group. Most dentate gyrus cells from control subjects have no inclusions, while FTLD, ALS, and AD subjects have significantly more granules per cell across the range of observed values. (G) The relationship between RBM45 nuclear inclusions and SAFB immunoreactivity is shown, along with the regression line. Regardless of disease state, with increasing numbers of RBM45 nuclear inclusions, SAFB immunoreactivity decreases, particularly at ≥ 3 granules per cell. The relationship between these two variables is statistically significant (*p* < 1 × 10^-16^). In the scatterplot shown, jitter is added to the points to more clearly visualize the data, but the separation between inclusion integer numbers is still seen. The inset image shows an example of adjacent cells in an FTLD subject where one cell with no RBM45 granules has strong SAFB nuclear immunoreactivity, while the neighboring cell has several nuclear RBM45 inclusions and low SAFB nuclear immunoreactivity. In (A-D), the scale bar = 20 μm, while in the inset in (G) the scale bar = 5 μm.

We focused our efforts on the dentate gyrus as RBM45 nuclear inclusions are significantly more prevalent in the dentate gyrus than in hippocampal pyramidal neurons^1^. Subject demographics are shown in Table 1. We used an image analysis pipeline to extract the number of cells, the number of RBM45 nuclear inclusions, and SAFB immunoreactivity from three dimensional images obtained via confocal microscopy (see Methods). Cytoplasmic RBM45 inclusions were counted manually. We counted a total of 8 fields per subject, corresponding to a total of 10,640 cells across all groups. In non-neurologic disease controls, RBM45 exhibited diffuse nuclear and strong perinucleolar immunoreactivity in dentate gyrus granule cells (Figure 7A, arrowheads). These cells also exhibited strong, diffuse nuclear SAFB immunoreactivity. In contrast, granule cells in FTLD, ALS, and AD patients typically exhibited multiple RBM45 nuclear inclusions with a corresponding loss of nuclear/perinucleolar RBM45 immunoreactivity (Figure 7B-D, respectively). RBM45 nuclear inclusions were negative for SAFB, consistent with our *in vitro* results. Cytoplasmic RBM45 inclusions were occasionally observed in dentate gyrus granule cells and these were positive for TDP-43 (Figure 7B-D, bottom panels [arrows]), but negative for SAFB.

We quantified the number of nuclear and cytoplasmic RBM45 inclusions in dentate gyrus granule cells from each subject. The distribution of nuclear inclusions exhibited a strong rightward skew, with many cells containing zero inclusions. Accordingly, we used the non-parametric Mann-Whitney U test to assess the differences in RBM45 puncta counts between groups. FTLD subjects had significantly more RBM45 inclusions than non-neurologic disease controls, AD, and ALS subjects (Figure 7E; *p* < 1 × 10^-6^ for control and AD and *p* < 0.01 for ALS versus FTLD, respectively). ALS subjects had significantly more nuclear inclusions than control and AD subjects (*p* < 1 × 10^-6^ and *p* < 0.01, respectively). Because non-neurologic disease controls had significantly more hippocampal dentate gyrus cells than FTLD, ALS, and AD subjects, we repeated this comparison after normalizing RBM45 inclusions to the number of cells and found that FTLD had significantly more nuclear inclusions per cell than ALS, AD, and control subjects (*p* < 1 × 10^-6^, *p* < 1 × 10^-8^, respectively). ALS subjects had significantly more RBM45 nuclear inclusions than control and AD subjects (*p* < 1 × 10^-6^ and *p* < 0.01, respectively). The comparison between AD and non-neurologic disease controls did not reach statistical significance following multiple testing correction (*p* = 0.015). We observed that cells containing RBM45 nuclear inclusions often had multiple inclusions (in some instances more than 10 per nucleus), irrespective of disease state (Figure 7F). The proportion of hippocampal cells containing at least 1 nuclear RBM45 inclusion was significantly greater than the proportion of cells containing at least 1 cytoplasmic inclusion (*p* < 1 × 10^-8^). The distribution of RBM45 nuclear inclusions per cell shows that the greatest proportion of positive cells and the highest numbers of inclusions per cell are found in the dentate gyrus of FTLD subjects (Figure 7E, F). AD and ALS subjects show comparable proportions and numbers of inclusions per cell, while the vast majority of cells from control subjects (> 85%) show 1 or fewer nuclear inclusions. Cytoplasmic RBM45 inclusions were not observed in non-neurologic disease controls. Summary statistics for RBM45 nuclear and cytoplasmic pathology for our analysis of dentate gyrus are shown in Table 2.

**Table 2.**
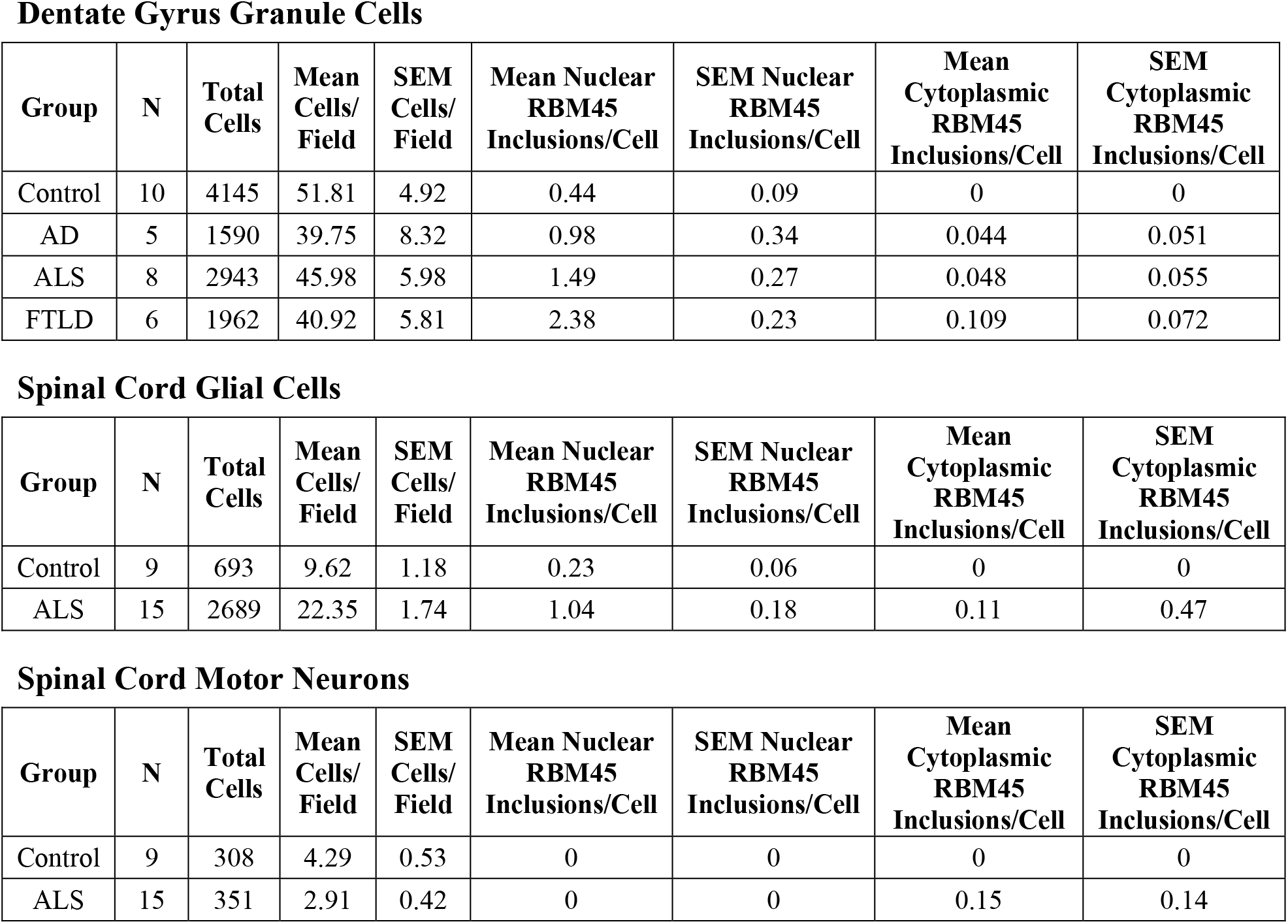
Summary Statistics from Tissue Immunohistochemistry Image Analysis. The table shows the number of individuals and cells counted for each tissue region and subject group. The number of neurons or glial cells and RBM45 nuclear and cytoplasmic inclusions were counted as described in the Methods section. For each subject, a total of 8 fields per region (hippocampal dentate gyrus or lumbar spinal cord) were evaluated.

Dentate Gyrus Granule Cells
| Group | N | Total Cells | Mean Cells/Field | SEM Cells/Field | Mean Nuclear RBM45 Inclusions/Cell | SEM Nuclear RBM45 Inclusions/Cell | Mean Cytoplasmic RBM45 Inclusions/Cell | SEM Cytoplasmic RBM45 Inclusions/Cell |
| --- | --- | --- | --- | --- | --- | --- | --- | --- |
| Control | 10 | 4145 | 51.81 | 4.92 | 0.44 | 0.09 | 0 | 0 |
| AD | 5 | 1590 | 39.75 | 8.32 | 0.98 | 0.34 | 0.044 | 0.051 |
| ALS | 8 | 2943 | 45.98 | 5.98 | 1.49 | 0.27 | 0.048 | 0.055 |
| FTLD | 6 | 1962 | 40.92 | 5.81 | 2.38 | 0.23 | 0.109 | 0.072 |

Spinal Cord Glial Cells
| Group | N | Total Cells | Mean Cells/Field | SEM Cells/Field | Mean Nuclear RBM45 Inclusions/Cell | SEM Nuclear RBM45 Inclusions/Cell | Mean Cytoplasmic RBM45 Inclusions/Cell | SEM Cytoplasmic RBM45 Inclusions/Cell |
| --- | --- | --- | --- | --- | --- | --- | --- | --- |
| Control | 9 | 693 | 9.62 | 1.18 | 0.23 | 0.06 | 0 | 0 |
| ALS | 15 | 2689 | 22.35 | 1.74 | 1.04 | 0.18 | 0.11 | 0.47 |

Table 2. Summary Statistics from Tissue Immunohistochemistry Image Analysis.
| Group | N | Total Cells | Mean Cells/Field | SEM Cells/Field | Mean Nuclear RBM45 Inclusions/Cell | SEM Nuclear RBM45 Inclusions/Cell | Mean Cytoplasmic RBM45 Inclusions/Cell | SEM Cytoplasmic RBM45 Inclusions/Cell |
| --- | --- | --- | --- | --- | --- | --- | --- | --- |
| Control | 9 | 308 | 4.29 | 0.53 | 0 | 0 | 0 | 0 |
| ALS | 15 | 351 | 2.91 | 0.42 | 0 | 0 | 0.15 | 0.14 |

In subjects with nuclear RBM45 inclusions, a bimodal pattern of SAFB immunoreactivity was often seen wherein cells with multiple RBM45 nuclear inclusions had low SAFB immunoreactivity and cells without RBM45 nuclear inclusions had high SAFB immunoreactivity. We quantified this relationship (Figure 7G) and used linear regression to assess its significance. Across all groups, increasing numbers of RBM45 nuclear inclusions per cell were associated with decreasing SAFB immunoreactivity (*p <* 1 × 10^-16^). SAFB nuclear immunoreactivity was lowest when cells had ≥ 3 RBM45 nuclear inclusions. This relationship could be observed within individual image fields from individual subjects (Figure 7G, inset).

We performed a similar analysis of lumbar spinal cord tissue sections from ALS and non-neurologic disease controls. In motor neurons from non-neurologic disease control subjects, RBM45 immunoreactivity was largely nucleolar (Figure 8A, arrowheads) and SAFB and TDP-43 immunoreactivity was diffuse. RBM45, TDP-43, or SAFB nuclear and cytoplasmic inclusion pathology was absent in control subjects (Figure 8A). In contrast, ALS patients frequently exhibited skein-like cytoplasmic RBM45 inclusion pathology that was also immunoreactive for TDP-43 but not SAFB (Figure 8B). We also noted a large decrease in RBM45 nucleolar immunoreactivity in spinal cord motor neurons with cytoplasmic RBM45 inclusions (Figure 8B, arrowheads). Despite altered RBM45 motor neuron nucleolar immunoreactivity, we did not identify any nuclear RBM45 inclusions in motor neurons in ALS subjects (Figure 8B, Table 2). Instead, we frequently observed RBM45 nuclear inclusions in glial cells adjacent to motor neurons in ALS, but not in non-neurologic disease controls (Figure 8B, arrows, 8C; Table 2).

**Figure 8.**
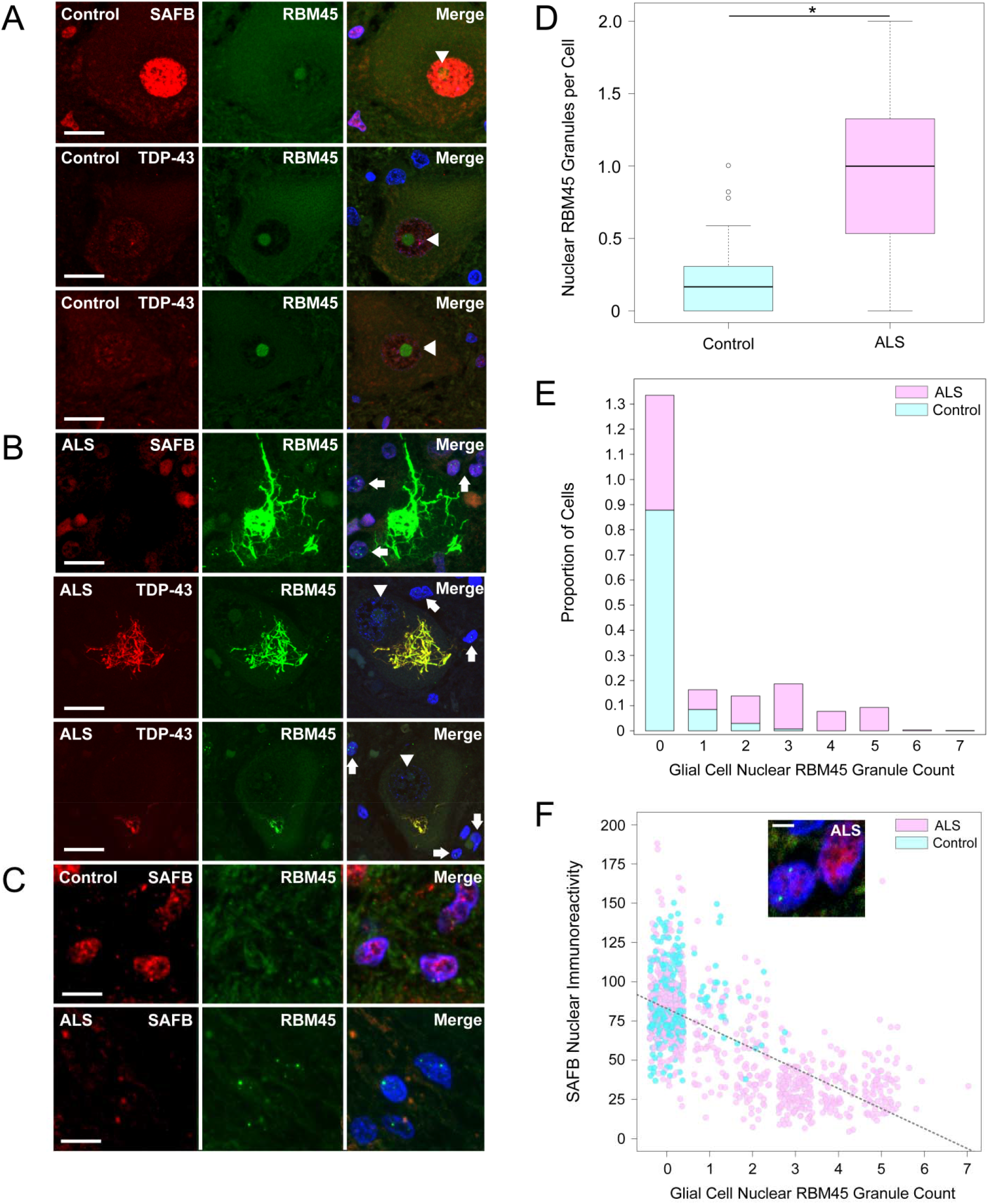
RBM45 Subcellular Distribution and Inclusion Pathology in Lumbar Spinal Cord. Immunohistochemistry for RBM45 with either SAFB or TDP-43 performed in lumbar spinal cord sections from ALS and non-neurologic disease control subjects. (A) RBM45, SAFB, and TDP-43 immunoreactivity were measured in motor neurons and glial cells of non-neurologic disease control subjects. RBM45 exhibited strong nucleoloar immunoreactivity in motor neurons (arrowheads), while SAFB and TDP-43 immunoreactivity were diffusely nuclear. (B) In ALS patients, motor neuron nucleolar RBM45 immunoreactivity was reduced (arrowheads) and large, skein-like cytoplasmic inclusions were commonly observed. These inclusions were positive for TDP-43 but negative for SAFB. ALS patient glial cells frequently contained multiple RBM45 nuclear inclusions (arrows). (C) Glial cells from non-neurologic disease control (top) and ALS (bottom) subjects. Glial cells from ALS subjects frequently contained 1 or more nuclear RBM45 inclusions. The presence of glial RBM45 nuclear inclusions was associated with reduced SAFB immunoreactivity. (D) Boxplot showing the number of glial nuclear RBM45 inclusions in control and ALS subjects. For the comparison shown, * = p < 1 × 10^-6^. (E) Stacked bar plot showing the proportion of cells (Y axis) with the given number glial nuclear RBM45 inclusions (X axis) across groups. ALS subjects have significantly more inclusions per glial cell than control subjects, who rarely exhibit RBM45 glial cell pathology. (F) The relationship between glial RBM45 nuclear inclusions and SAFB immunoreactivity is shown, along with the regression line. Increasing numbers of RBM45 nuclear inclusions are associated with reduced SAFB immunoreactivity, particularly at ≥ 3 granules per cell. The relationship between these two variables is statistically significant (*p* < 1 × 10^-6^). In the scatterplot shown, jitter is added to the points to more clearly visualize the data, but the separation between inclusion integer numbers can still be seen. The inset image shows an example of adjacent glial cells in an ALS patient where one cell without RBM45 nuclear inclusions has strong SAFB nuclear immunoreactivity, while the neighboring cell with RBM45 nuclear inclusions shows reduced SAFB immunoreactivity. In (A and B) the scale bar = 20 μm, in (C) the scale bar = 5 μm, and for the inset in (F) the scale bar = 2.5 μm.

We then counted lumbar spinal cord glial cells and glial nuclear RBM45 inclusions in individual image fields. A total of 3,382 lumbar spinal cord glial cells across 26 subjects were counted using this approach. Glial cells in ALS patients contained significantly more RBM45 nuclear inclusions per cell than those from non-neurologic disease controls (Figure 8D, Mann-Whitney *p <* 1 × 10^-8^). We also compared the proportion of spinal cord glial cells with nuclear RBM45 inclusions and cytoplasmic inclusions. We found that the proportion of cells with at least 1 RBM45 nuclear inclusion was significantly greater than the proportion of glial cells with at least 1 RBM45 cytoplasmic inclusion (Figure 8E, *p* < 1 × 10^-5^). As in our analysis of hippocampal tissue, we found that the majority of glial cells in non-neurologic disease controls (≥ 85%) lack RBM45 nuclear inclusions (Figure 8E) and did not contain cytoplasmic RBM45 inclusions. Likewise, glial cells that contained multiple RBM45 nuclear inclusions showed reduced SAFB immunoreactivity (Figure 8F, inset). We explored the relationship between nuclear RBM45 inclusion number and SAFB immunoreactivity and found that greater numbers of RBM45 inclusions per cell were associated with decreasing SAFB nuclear immunoreactivity (Figure 8F, *p* < 1 × 10^-6^). SAFB nuclear immunoreactivity was lowest when cells had ≥ 3 RBM45 nuclear inclusions (Figure 8F). Summary statistics for the image analysis of human spinal cord tissue are presented in Table 2. Collectively, our results indicate that nuclear RBM45 pathology is a common occurrence that exhibits cell-type specific patterns in FTLD, AD, and ALS. Moreover, nuclear RBM45 inclusions are found in significantly higher proportions of cells than cytoplasmic RBM45 inclusions in FTLD, ALS, and AD. Finally, RBM45 nuclear inclusions are devoid of the NSB marker protein SAFB, consistent with our in vitro results during chronic stress (Figures 4, 6).

## Discussion

The goals of this study were to understand whether RBM45 associates with nuclear organelles, whether this association promotes the aggregation and formation of nuclear RBM45 inclusions, and to understand the cell type- and disease-specific specific patterns of RBM45 immunoreactivity and inclusion pathology in FTLD, ALS, and AD. We found that RBM45 associates with nuclear stress bodies (NSBs), stress-induced RNA-protein complexes, and that the persistent association of RBM45 in NSBs is sufficient to produce insoluble nuclear RBM45 inclusions. This association is mediated RNA recognition motifs (RRM) 2 and 3 in RBM45. In addition, the chronic sequestration of RBM45 in NSBs was sufficient to promote inclusion formation, even when other NSB marker proteins had disassembled from these complexes. In human CNS tissue, nuclear RBM45 inclusions were found in ALS, FTLD, and AD in distinct cell types and this pathology occurs more frequently than cytoplasmic RBM45 pathology.

Aggregation and assembly into membraneless organelles is essential to many RBP functions, such as the regulation of transcription, as well as mRNA splicing, transport, and decay^53^. The assembly of RBPs, nucleic acids, and other factors into membraneless organelles acts to compartmentalize these components, leading to a high local concentration of enzymes and substrates of the associated biochemical reactions^53–55^. While RBP association and oligomerization may, therefore, be crucial to the normal functions of these proteins, evidence also points to oligomerization as a key driver of inclusion formation in ALS and FTLD^13, 56^. Our prior work demonstrated that RBM45 forms oligomeric complexes and interacts with many other RBPs via an intrinsically disordered peptide segment termed the homo-oligomer assembly (HOA) domain^3^. We sought to understand whether these properties also lead to the association of RBM45 with a nuclear organelle. We examined the co-localization of RBM45 and marker proteins of several known membraneless, RBP-containing organelles, including nuclear speckles, Cajal bodies, nuclear gems, nuclear stress bodies (NSBs), and cytoplasmic stress granules. Our results indicate that under basal conditions RBM45 does not co-localize with any of these organelles and, instead, exhibits a diffuse nuclear localization. Following the onset of cellular stress, however, RBM45 is incorporated into NSBs.

NSBs are protein-RNA complexes that form in response to stress-induced transcription of satellite III (SatIII) repeats from pericentromeric heterochromatin^50^. The resultant SatIII transcripts act as scaffolds that recruit various RBPs, notably the transcription factor HSF1 and the hnRNP SAFB, to NSBs, resulting in the appearance of several nuclear granules that dissipate following stressor removal^42, 50^. Despite a well-characterized mechanism of formation, the functions of NSBs have remained enigmatic. Current theory suggests that NSBs act as one component of a larger gene expression regulatory program for the cellular response to stress^50^. By sequestering various DNA/RNA binding proteins, such as RBM45, SAFB, and HSF1, NSBs are thought to shift gene expression in ways that alter both transcription and splicing events. Further studies are needed to define the role of RBM45 on NSB function(s) and how incorporation into NSBs alters the gene regulatory functions of RBM45. A key insight from this work is that the persistent association of RBM45 with NSBs is sufficient to promote the protein’s aggregation into inclusions (Figures 4, 5, and 6), which may further impact the normal function of RBM45 and other proteins contained within these inclusions.

RBPs can be recruited to NSBs by direct binding to SatIII transcripts or indirectly via protein-protein interactions (PPI) with resident NSB proteins^50^. Our data indicates that recruitment of RBM45 to NSBs requires the RNA binding motifs (RRM2 and RRM3) and does not require the HOA domain important for PPI with other RBPs (Figure 3). This suggests that RBM45 associates with NSBs via direct RNA binding. In our previous efforts to characterize RBM45 PPIs, we did not identify NSB proteins HSF1 and SAFB (the canonical NSB marker proteins), which also do not physically interact^5^. Prior studies demonstrated that RBM45 has a binding preference for poly(C) and poly(G) RNA^6^. The SatIII transcripts that act as the scaffolds for NSBs are themselves G- and C-rich^41, 42^, consistent with our hypothesis that RBM45 incorporation into NSBs is a consequence of direct binding to SatIII transcripts. We observed that RBM45 is not required for the formation of NSBs (Figure S4). This result is not unexpected given that the mechanism of NSB formation is transcription of SatIII DNA by HSF1 binding rapidly following stress onset^42, 50^. NSB formation also exhibits cell- and stressor-specific variability and even HSF1 itself is dispensable for NSB formation during certain types of cellular stress^31^. Further studies are required to determine the mechanisms by which constituent proteins are recruited to NSBs.

RBM45 is frequently observed in cytoplasmic and nuclear inclusions in neurons and glia in FTLD, ALS, and AD, but not in non-neurologic disease controls^1, 2^. This suggests that ongoing cellular stressors or pathological processes alter the normal diffuse nuclear localization of RBM45. RBM45 interacts with both TDP-43 and FUS^3^, is found in TDP-43-positive neuronal cytoplasmic inclusions in ALS, FTLD, and AD, and has previously been reported to associate with cytoplasmic SGs^3, 4^. We examined the distribution of RBM45 during conditions of cellular stress and found no evidence of RBM45 cytoplasmic translocation or association with SGs (Figure 1D-E and Figure 2). Several factors may account for the conflicting claims regarding RBM45 and SGs made here and elsewhere. First, previous claims of RBM45 association with SGs were made using plasmid-based overexpression of RBM45 with a mutated, non-functional NLS. While such approaches establish that RBM45 can associate with SGs, they rely on removing RBM45 from its normal subcellular milieu and expressing the protein at non-physiological levels. Our study using wild-type RBM45 expression at physiologic levels failed to identify RBM45 association with cytoplasmic SGs under a diverse array of stress conditions. Consistent with this observation, a recent study to comprehensively characterize the SG proteome by mass spectrometry failed to detect RBM45 within SGs^57^.

Moreover, species- and cell type-specific differences in NSB and RBM45 biochemistry may be crucial determinants of a cell’s potential for RBM45 nuclear aggregation and inclusion formation. Notably, NSBs are specific to primate species and are not observed in rodent cells^42, 50^. Thus, the stress-associated functions of RBM45 are likely to differ across species. Similarly, RBM45 exhibits a predominantly nucleolar immunoreactivity in motor neurons in human CNS tissue that was not seen in any other cell type examined (Figure 8). Lumbar spinal cord motor neurons also did not exhibit nuclear RBM45 inclusions (Figure 8), raising the intriguing possibility that RBM45 does not associate with NSBs in some cell populations and that these cell types do not develop RBM45 nuclear inclusion pathology as a result. Additional studies are required to define the distribution of RBM45 protein in various cell types and the factors that regulate the subcellular localization of RBM45. One outstanding question is how the predominantly nuclear RBM45 becomes incorporated into cytoplasmic, TDP-43 positive inclusions in FTLD, ALS, and AD (Figures 7, 8). While RBM45 is a predominantly nuclear protein, low amounts of cytoplasmic RBM45 are detected by Western blot^3^. Since RBM45 and TDP-43 interact, the association of cytoplasmic RBM45 with TDP-43 may be sufficient to result in RBM45 incorporation into cytoplasmic inclusions observed in ALS, FTLD, and AD. Moreover, the nucleocytoplasmic transport defects seen in ALS/FTLD may result in accumulation of cytoplasmic RBM45, which may promote its incorporation into inclusion bodies in these diseases^58, 59^.

RBM45 assembles into higher-order oligomers^3^ and this behavior is likely essential to one or more of the protein’s functions, similar to other aggregation-prone RBPs. While acute stress results in transient RBM45 association with NSBs, chronic, low levels of cellular stress were sufficient to trap RBM45 in large nuclear aggregates. Importantly, in the absence of SatIII transcripts, this phenomenon was not observed (Figure S3), suggesting that RBM45 nuclear aggregation and inclusion formation requires the initial formation of RBM45 containing NSBs. RBM45 nuclear aggregates that result from chronic stress were typically negative for other NSB marker proteins, despite initial co-localization in NSBs (Figure 4). Similarly, we did not observe co-localization of RBM45 and NSB marker proteins in ALS, FTLD, or AD tissue (Figures 7 and 8). Several factors may account for these observations. First, the HOA domain that mediates RBM45 self-association is an intrinsically disordered peptide segment that promotes the assembly of RBM45 oligomers^3^. Chronic stress likely facilitates this process via compartmentalization in NSBs in a manner analogous to the sequestration and aggregation of TDP-43 and FUS in cytoplasmic SGs. Second, HSF1 and SAFB do not oligomerize into large, macromolecular assemblies as part of their normal functions. HSF1 functions as a DNA binding-competent trimer and is only detectable in NSBs over a limited time course, specifically the early, but not late, phase of NSB lifetime^50^. SAFB likewise functions as a monomer or homodimer, but has not previously been shown to assemble into oligomeric complexes. It too, is detected over a limited time frame in the lifespan of a NSB, appearing approximately 1 hour after HSF1 granules are detectable^50^. Despite the fact that RBM45, HSF1, and SAFB are all incorporated into NSBs, no direct physical interaction between these proteins has previously been observed^5^. The physical and temporal separation of these proteins, together with the low aggregation potential of HSF1 and SAFB, provide plausible explanations for the absence of these proteins in RBM45 nuclear inclusions resulting from chronic stress. RBM45 nuclear inclusions may nonetheless contain additional proteins and future studies will explore the proteome of these structures.

To understand cell type- and disease-specific patterns of RBM45 inclusion pathology in neurodegenerative diseases, we performed an extensive quantification of RBM45 pathology in ALS, FTLD, AD, and non-neurologic disease controls. In hippocampal dentate gyrus neurons, numerous RBM45 nuclear inclusions with a punctate morphology were observed in FTLD, AD, and ALS subjects, but not in non-neurologic disease controls. When we quantified the number of cells and RBM45 nuclear inclusions in each cell, we found that FTLD subjects had (1) the highest proportion of cells with at least 1 nuclear inclusion, (2) the highest average number of nuclear inclusions per cell, and (3) the highest number of nuclear inclusions observed within a single cell. RBM45 nuclear inclusions were not positive for TDP-43 or SAFB. Instead, we found that the presence of RBM45 nuclear inclusions correlated with disruption of the normal distribution and immunoreactivity of SAFB in ALS, FTLD, and AD (Figures 7 and 8). SAFB immunoreactivity was lowest when cells had 3 or more RBM45 nuclear inclusions (Figure 7D). Critically, this was not due to a general decrease in SAFB immunoreactivity in disease, as cells without RBM45 nuclear inclusions showed strong SAFB immunoreactivity comparable to that in non-neurologic disease controls, even when these cells were adjacent to cells with RBM45 nuclear inclusions (Figures 7G and 8F, inset). The distribution of RBM45 nuclear inclusions in dentate gyrus granule cells ranged from 0 to 17, with the majority of cells harboring > 5 inclusions found in FTLD subjects. This range of values shows good agreement with previous efforts to quantify the number of NSBs that form in response to different stressors in cultured cells. One study found that individual HeLa cells typically produced 1-20 NSBs in response to a variety of stressors, with 50% of cells harboring 1-5 NSBs^31^. Our results suggest the RMB45 nuclear inclusions may denote the prior generation of NSBs in these cells. In some instances, we observed RBM45 nuclear inclusions in non-neurologic disease controls, though this may be due to the normal aging process as seen with other RBP pathology in aged non-neurologic disease CNS tissues^60, 61^.

We also quantified RBM45 pathology in the lumbar spinal cord of ALS and non-neurologic disease controls. As in our prior study, we found TDP-43-positive RBM45 cytoplasmic inclusions in motor neurons of ALS but not in non-neurologic control subjects (Figure 8). A key finding from this study is RBM45 nuclear inclusions show cell-type specificity in ALS spinal cord. Specifically, we observed RBM45 nuclear inclusions in ALS glial cells, but not motor neurons. We hypothesize that the RBM45 nucleolar localization in motor neurons limits the protein’s ability to associate with NSBs, and hence, form nuclear inclusions. Alternatively, spinal cord motor neurons may not form NSBs upon cellular stress and therefore do not subsequently form RBM45 nuclear inclusions. We did observe that cytoplasmic sequestration of RBM45 in inclusions is associated with reduced nucleolar RBM45 immunoreactivity (Figure 8). In contrast to motor neurons, glial cells in the lumbar spinal cord of ALS subjects frequently had RBM45 nuclear inclusions (Figure 8). Notably, the proportion of glial cells with at least one RBM45 nuclear inclusion was significantly greater than the proportion of glial cells with cytoplasmic RBM45 inclusions (Figure 8 and Table 2). Similar to our results from the dentate gyrus, we also found that glial SAFB immunoreactivity was reduced in the presence of RBM45 nuclear inclusions. Glial inclusion pathology occurs in various forms of ALS and glial inclusions may contain TDP-43, FUS, SOD1, C9ORF72-associated dipeptide repeat proteins (DPRs), or other ALS-associated proteins^62, 63^. The occurrence of RBM45 inclusions in both the cytoplasm and nucleus of ALS glial cells, together with the diversity of other proteins found in glial inclusions in ALS, underscores the predominant role of glial dysfunction in the pathobiology of ALS and related diseases.

Overall, our results demonstrate that RBM45 associates with NSBs during cellular stress and that chronic stress leads to persistent self-association of RBM45 and nuclear RBM45 inclusion formation. Nuclear RBM45 inclusions are extensively found in hippocampal dentate gyrus cells in ALS, FTLD, and AD, as well as in glial cells in the spinal cord of ALS patients. The presence of nuclear inclusions is associated with depletion of RBM45 from its normal subcellular milieu and altered distribution of other RBPs. Collectively, our results add new insights into the biological functions of RBM45, the mechanisms by which RBM45 forms inclusions, and implicate NSBs in the pathogenesis of several neurodegenerative diseases.

## Acknowledgements

This work was supported by National Institutes of Health grants NS068179 to RB and NS080614 to MC, the Achievement Rewards for College Scientists Foundation, Inc. Pittsburgh Chapter scholarship to MC, and support from the Barrow Neurological Foundation to RB. The authors wish to thank the Target ALS Multicenter Postmortem Tissue Core for human tissue samples used in this study.

---

**Supplementary Figure 1.**
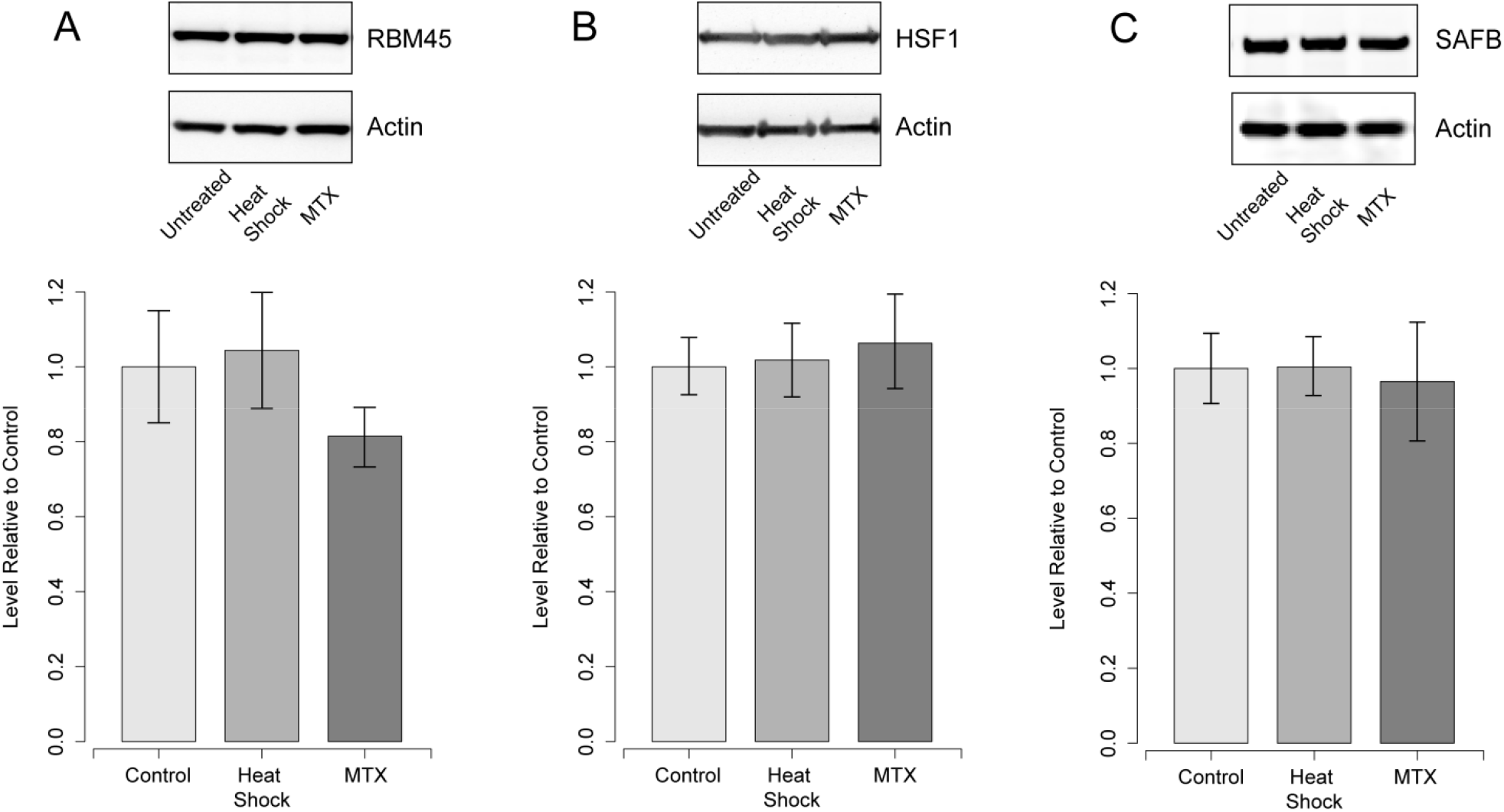
NSB Protein Levels following Cellular Stress. Total protein extracts were prepared from untreated cells, heat shocked cells (42°C for 2 hours), and cells treated with the genotoxic stressor mitoxantrone (MTX; 20 µM for 6 hours). Each panel shows the results of loading 10 μg of each extract and blotting for the indicated proteins with actin used as a loading control. (A) RBM45; (B) HSF1; (C) SAFB. No statistically significant differences in the levels of the indicated proteins were detected between conditions (*p >* 0.05).

**Supplementary Figure 2.**
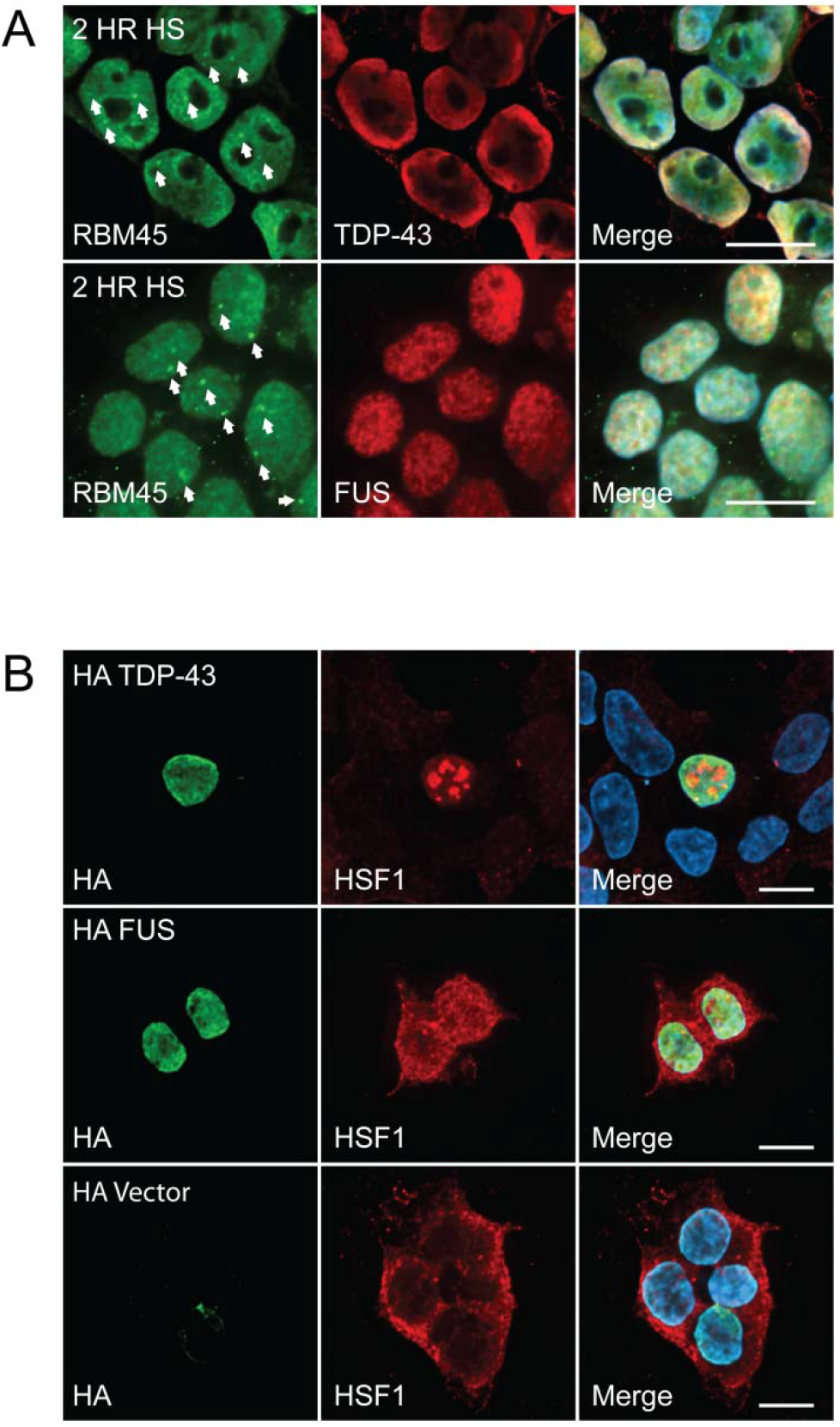
Assessment of the Association of TDP-43 and FUS with NSBs. (A) HEK293 cells were heat-shocked for 2 hours at 42°C to induce formation of NSBS. Cells were stained for RBM45 and TDP43 or FUS as indicated. Following heat shock, numerous RBM45-positive NSBs become visible in the cell nucleus (arrows) and these do not contain TDP-43 or FUS. Some TDP-43-positive stress granules are visible in the cytoplasm following heat shock and these do not contain RBM45. (B) HEK293 cells were transiently transfected to overexpress HA-tagged TDP-43 or FUS and were then stained for the HA tag and the NSB marker HSF1. Overexpression of TDP-43 was sufficient to robustly induce NSB formation, but NSBs were negative for TDP-43. Neither overexpression of FUS or transfection with a control HA vector resulted in NSB formation. For all images, scale bar = 10 μm.

**Supplementary Figure 3.**
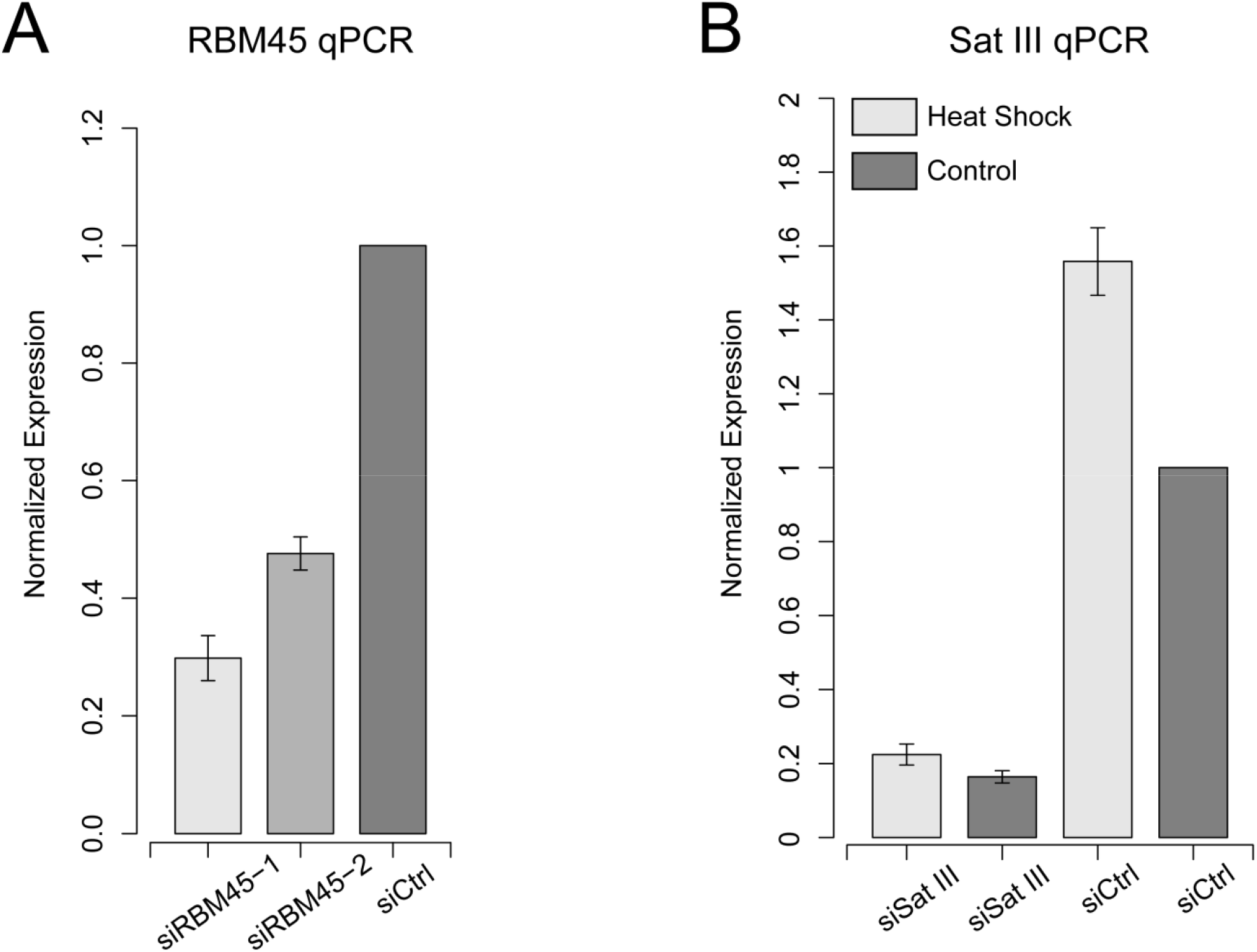
siRNA Efficiency. HEK293 cells were transfected with the indicated siRNA and transcript levels were measured by real-time PCR. Each bar presents the relative transcript abundance, expressed as a proportion of the corresponding transcript level in untreated cells transfected with a scrambled control siRNA. (A) Evaluation of two unique siRNAs, each targeting RBM45. (B) Evaluation of SatIII knockdown efficiency in untreated and heat shocked (42°C for 2 hours) cells.

**Supplementary Figure 4.**
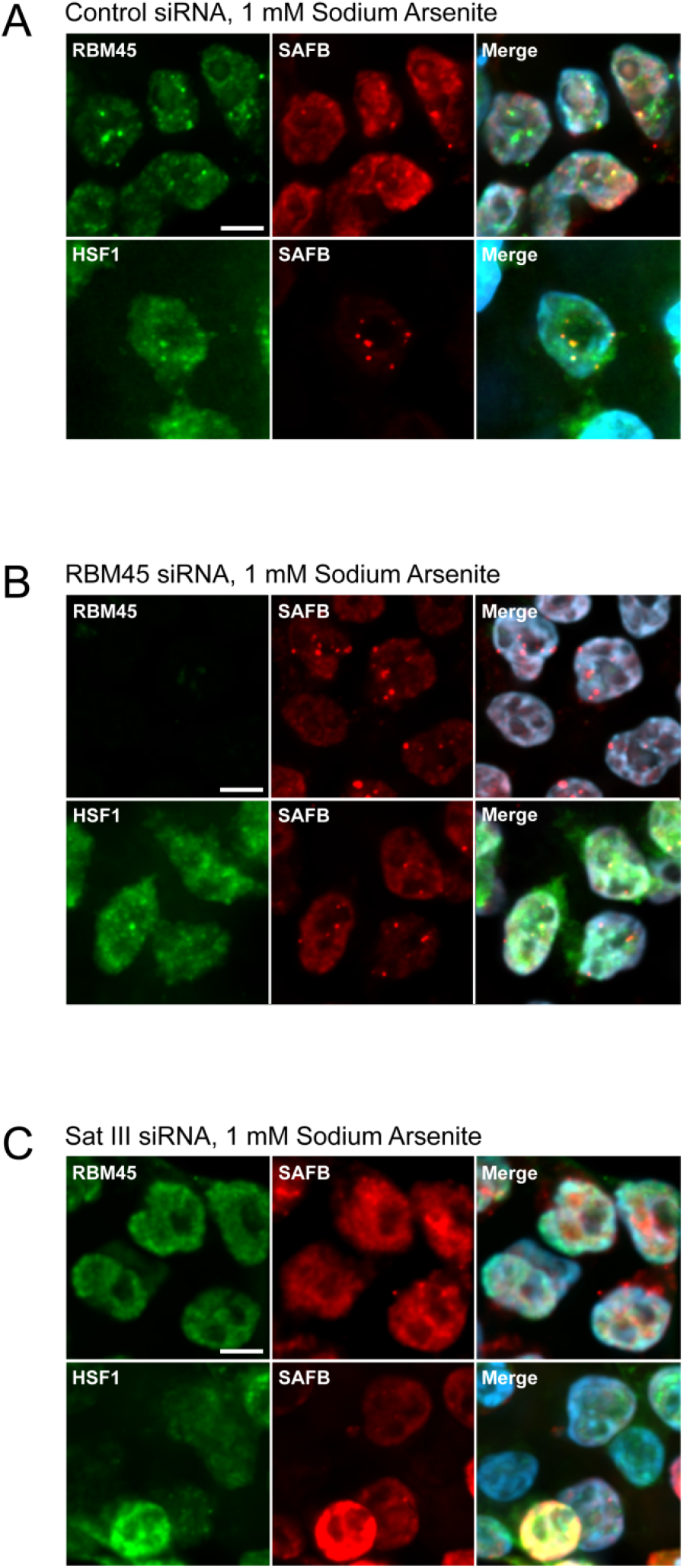
Effect of siRNA knockdown of RBM45 and SatIII on NSB formation. HEK293 cells were transfected with siRNAs targeting RBM45, SatIII, or off-target scrambled siRNAs (control). Nuclear stress body (NSB) formation was then assessed by immunocytochemistry. Cells were treated with 1 mM sodium arsenite for 1 hour to induce NSB formation. (A) Effect of off-target, scrambled siRNA (control) on NSB formation. Cells transfected with control siRNAs readily form SAFB, RBM45, and HSF1-positive nuclear stress bodies following treatment with sodium arsenite. (B) Cells transfected with siRNAs targeting RBM45 show reduced levels of RBM45, but readily form SAFB and HSF1-positive NSBs following cellular stress. (C) Effect of SatIII knockdown on NSB formation. Knockdown of SatIII leads to a loss of NSB formation during cellular stress as indicated by the loss of SAFB, RBM45, and HSF1-positive NSBs in cells transfected with SatIII targeting siRNAs. For all images, scale bar = 5 μm.

**Supplementary Figure 5.**
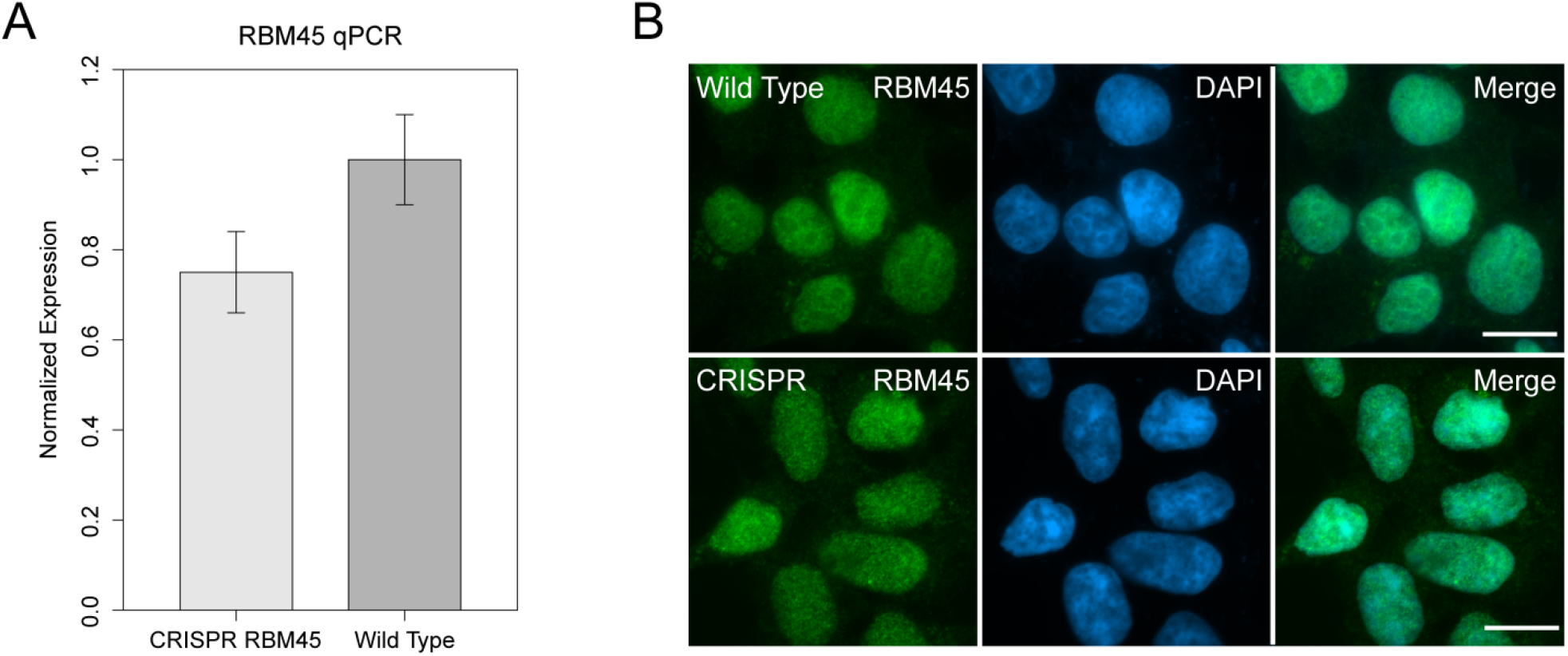
Characterization of CRISPR-edited FLAG-tagged RBM45 Expressing Cells. CRISPR-cas9 genome editing was used to generate HEK293 cells expressing N-terminally 2x FLAG-tagged RBM45. (A) RBM45 transcript levels were evaluated by real-time PCR. The barplot presents the mean RBM45 transcript levels relative to unedited cells expressing wild-type RBM45. (B) Immunofluorescence was performed using an antibody to RBM45 in CRISPR-edited HEK293 cells. The results show that the abundance and subcellular distribution of RBM45 protein and HEK293 cell nuclear morphology are not altered by the 2X FLAG tag.

## References

1. Collins, M.; Riascos, D.; Kovalik, T.; An, J.; Krupa, K.; Krupa, K.; Hood, B. L.; Conrads, T. P.; Renton, A. E.; Traynor, B. J.; Bowser, R., The RNA-binding motif 45 (RBM45) protein accumulates in inclusion bodies in amyotrophic lateral sclerosis (ALS) and frontotemporal lobar degeneration with TDP-43 inclusions (FTLD-TDP) patients. Acta Neuropathol 2012, 124 (5), 717–32.

2. Konno, T.; Tada, M.; Shiga, A.; Tsujino, A.; Eguchi, H.; Masuda-Suzukake, M.; Hasegawa, M.; Nishizawa, M.; Onodera, O.; Kakita, A.; Takahashi, H., C9ORF72 repeat-associated non-ATG-translated polypeptides are distributed independently of TDP-43 in a Japanese patient with c9ALS. Neuropathol Appl Neurobiol 2014, 40 (6), 783–8.

3. Li, Y.; Collins, M.; Geiser, R.; Bakkar, N.; Riascos, D.; Bowser, R., RBM45 homo-oligomerization mediates association with ALS-linked proteins and stress granules. Sci Rep 2015, 5, 14262.

4. Mashiko, T.; Sakashita, E.; Kasashima, K.; Tominaga, K.; Kuroiwa, K.; Nozaki, Y.; Matsuura, T.; Hamamoto, T.; Endo, H., Developmentally Regulated RNA-binding Protein 1 (Drb1)/RNA-binding Motif Protein 45 (RBM45), a Nuclear-Cytoplasmic Trafficking Protein, Forms TAR DNA-binding Protein 43 (TDP-43)-mediated Cytoplasmic Aggregates. J Biol Chem 2016, 291 (29), 14996-5007.

5. Li, Y.; Collins, M.; An, J.; Geiser, R.; Tegeler, T.; Tsantilas, K.; Garcia, K.; Pirrotte, P.; Bowser, R., Immunoprecipitation and mass spectrometry defines an extensive RBM45 protein-protein interaction network. Brain Res 2016, 1647, 79–93.

6. Tamada, H.; Sakashita, E.; Shimazaki, K.; Ueno, E.; Hamamoto, T.; Kagawa, Y.; Endo, H., cDNA cloning and characterization of Drb1, a new member of RRM-type neural RNA-binding protein. Biochem Biophys Res Commun 2002, 297 (1), 96–104.

7. Uversky, V. N., The roles of intrinsic disorder-based liquid-liquid phase transitions in the “Dr. Jekyll-Mr. Hyde” behavior of proteins involved in amyotrophic lateral sclerosis and frontotemporal lobar degeneration. Autophagy 2017, 13 (12), 2115–2162.

8. Harrison, A. F.; Shorter, J., RNA-binding proteins with prion-like domains in health and disease. Biochem J 2017, 474 (8), 1417–1438.

9. Zhu, L.; Brangwynne, C. P., Nuclear bodies: the emerging biophysics of nucleoplasmic phases. Curr Opin Cell Biol 2015, 34, 23–30.

10. Maziuk, B.; Ballance, H. I.; Wolozin, B., Dysregulation of RNA Binding Protein Aggregation in Neurodegenerative Disorders. Front Mol Neurosci 2017, 10, 89.

11. Mackenzie, I. R.; Nicholson, A. M.; Sarkar, M.; Messing, J.; Purice, M. D.; Pottier, C.; Annu, K.; Baker, M.; Perkerson, R. B.; Kurti, A.; Matchett, B. J.; Mittag, T.; Temirov, J.; Hsiung, G. R.; Krieger, C.; Murray, M. E.; Kato, M.; Fryer, J. D.; Petrucelli, L.; Zinman, L.; Weintraub, S.; Mesulam, M.; Keith, J.; Zivkovic, S. A.; Hirsch-Reinshagen, V.; Roos, R. P.; Zuchner, S.; Graff-Radford, N. R.; Petersen, R. C.; Caselli, R. J.; Wszolek, Z. K.; Finger, E.; Lippa, C.; Lacomis, D.; Stewart, H.; Dickson, D. W.; Kim, H. J.; Rogaeva, E.; Bigio, E.; Boylan, K. B.; Taylor, J. P.; Rademakers, R., TIA1 Mutations in Amyotrophic Lateral Sclerosis and Frontotemporal Dementia Promote Phase Separation and Alter Stress Granule Dynamics. Neuron 2017, 95 (4), 808–816 e9.

12. Conicella, A. E.; Zerze, G. H.; Mittal, J.; Fawzi, N. L., ALS Mutations Disrupt Phase Separation Mediated by alpha-Helical Structure in the TDP-43 Low-Complexity C-Terminal Domain. Structure 2016, 24 (9), 1537–49.

13. Murakami, T.; Qamar, S.; Lin, J. Q.; Schierle, G. S.; Rees, E.; Miyashita, A.; Costa, A. R.; Dodd, R. B.; Chan, F. T.; Michel, C. H.; Kronenberg-Versteeg, D.; Li, Y.; Yang, S. P.; Wakutani, Y.; Meadows, W.; Ferry, R. R.; Dong, L.; Tartaglia, G. G.; Favrin, G.; Lin, W. L.; Dickson, D. W.; Zhen, M.; Ron, D.; Schmitt-Ulms, G.; Fraser, P. E.; Shneider, N. A.; Holt, C.; Vendruscolo, M.; Kaminski, C. F.; St George-Hyslop, P., ALS/FTD Mutation-Induced Phase Transition of FUS Liquid Droplets and Reversible Hydrogels into Irreversible Hydrogels Impairs RNP Granule Function. Neuron 2015, 88 (4), 678–90.

14. Wolozin, B.; Ivanov, P., Stress granules and neurodegeneration. Nat Rev Neurosci 2019, 20 (11), 649–666.

15. Li, Y. R.; King, O. D.; Shorter, J.; Gitler, A. D., Stress granules as crucibles of ALS pathogenesis. J Cell Biol 2013, 201 (3), 361–72.

16. Anderson, P.; Kedersha, N., Stress granules: the Tao of RNA triage. Trends Biochem Sci 2008, 33 (3), 141–50.

17. Kedersha, N.; Anderson, P., Regulation of translation by stress granules and processing bodies. Prog Mol Biol Transl Sci 2009, 90, 155–85.

18. Vance, C.; Scotter, E. L.; Nishimura, A. L.; Troakes, C.; Mitchell, J. C.; Kathe, C.; Urwin, H.; Manser, C.; Miller, C. C.; Hortobagyi, T.; Dragunow, M.; Rogelj, B.; Shaw, C. E., ALS mutant FUS disrupts nuclear localization and sequesters wild-type FUS within cytoplasmic stress granules. Hum Mol Genet 2013, 22 (13), 2676–88.

19. Colombrita, C.; Zennaro, E.; Fallini, C.; Weber, M.; Sommacal, A.; Buratti, E.; Silani, V.; Ratti, A., TDP-43 is recruited to stress granules in conditions of oxidative insult. J Neurochem 2009, 111 (4), 1051–61.

20. Parker, S. J.; Meyerowitz, J.; James, J. L.; Liddell, J. R.; Crouch, P. J.; Kanninen, K. M.; White, A. R., Endogenous TDP-43 localized to stress granules can subsequently form protein aggregates. Neurochem Int 2012, 60 (4), 415–24.

21. Liu-Yesucevitz, L.; Bilgutay, A.; Zhang, Y. J.; Vanderweyde, T.; Citro, A.; Mehta, T.; Zaarur, N.; McKee, A.; Bowser, R.; Sherman, M.; Petrucelli, L.; Wolozin, B., Tar DNA binding protein-43 (TDP-43) associates with stress granules: analysis of cultured cells and pathological brain tissue. PLoS One 2010, 5 (10), e13250.

22. Bentmann, E.; Neumann, M.; Tahirovic, S.; Rodde, R.; Dormann, D.; Haass, C., Requirements for stress granule recruitment of fused in sarcoma (FUS) and TAR DNA-binding protein of 43 kDa (TDP-43). J Biol Chem 2012, 287 (27), 23079–94.

23. Johnson, B. S.; Snead, D.; Lee, J. J.; McCaffery, J. M.; Shorter, J.; Gitler, A. D., TDP-43 is intrinsically aggregation-prone, and amyotrophic lateral sclerosis-linked mutations accelerate aggregation and increase toxicity. J Biol Chem 2009, 284 (30), 20329–39.

24. Fushimi, K.; Long, C.; Jayaram, N.; Chen, X.; Li, L.; Wu, J. Y., Expression of human FUS/TLS in yeast leads to protein aggregation and cytotoxicity, recapitulating key features of FUS proteinopathy. Protein Cell 2011, 2 (2), 141–9.

25. Molliex, A.; Temirov, J.; Lee, J.; Coughlin, M.; Kanagaraj, A. P.; Kim, H. J.; Mittag, T.; Taylor, J. P., Phase separation by low complexity domains promotes stress granule assembly and drives pathological fibrillization. Cell 2015, 163 (1), 123–33.

26. Luo, F.; Gui, X.; Zhou, H.; Gu, J.; Li, Y.; Liu, X.; Zhao, M.; Li, D.; Li, X.; Liu, C., Atomic structures of FUS LC domain segments reveal bases for reversible amyloid fibril formation. Nat Struct Mol Biol 2018, 25 (4), 341–346.

27. Bowden, H. A.; Dormann, D., Altered mRNP granule dynamics in FTLD pathogenesis. J Neurochem 2016, 138 *Suppl 1*, 112–33.

28. Barmada, S. J.; Skibinski, G.; Korb, E.; Rao, E. J.; Wu, J. Y.; Finkbeiner, S., Cytoplasmic mislocalization of TDP-43 is toxic to neurons and enhanced by a mutation associated with familial amyotrophic lateral sclerosis. J Neurosci 2010, 30 (2), 639–49.

29. Polymenidou, M.; Lagier-Tourenne, C.; Hutt, K. R.; Bennett, C. F.; Cleveland, D. W.; Yeo, G. W., Misregulated RNA processing in amyotrophic lateral sclerosis. Brain Res 2012, 1462, 3–15.

30. Shelkovnikova, T. A.; Robinson, H. K.; Troakes, C.; Ninkina, N.; Buchman, V. L., Compromised paraspeckle formation as a pathogenic factor in FUSopathies. Hum Mol Genet 2014, 23 (9), 2298–312.

31. Valgardsdottir, R.; Chiodi, I.; Giordano, M.; Rossi, A.; Bazzini, S.; Ghigna, C.; Riva, S.; Biamonti, G., Transcription of Satellite III non-coding RNAs is a general stress response in human cells. Nucleic Acids Res 2008, 36 (2), 423–34.

32. Denegri, M.; Chiodi, I.; Corioni, M.; Cobianchi, F.; Riva, S.; Biamonti, G., Stress-induced nuclear bodies are sites of accumulation of pre-mRNA processing factors. Mol Biol Cell 2001, 12 (11), 3502–14.

33. Chiodi, I.; Biggiogera, M.; Denegri, M.; Corioni, M.; Weighardt, F.; Cobianchi, F.; Riva, S.; Biamonti, G., Structure and dynamics of hnRNP-labelled nuclear bodies induced by stress treatments. J Cell Sci 2000, 113 *(Pt* *22*), 4043–53.

34. Kolarcik, C. L.; Bowser, R., Retinoid signaling alterations in amyotrophic lateral sclerosis. Am J Neurodegener Dis 2012, 1 (2), 130–45.

35. Kedersha, N.; Anderson, P., Mammalian stress granules and processing bodies. Methods Enzymol 2007, 431, 61–81.

36. Busa, R.; Geremia, R.; Sette, C., Genotoxic stress causes the accumulation of the splicing regulator Sam68 in nuclear foci of transcriptionally active chromatin. Nucleic Acids Res 2010, 38 (9), 3005–18.

37. Aiyar, A.; Xiang, Y.; Leis, J., Site-directed mutagenesis using overlap extension PCR. Methods Mol Biol 1996, 57, 177–91.

38. Ran, F. A.; Hsu, P. D.; Wright, J.; Agarwala, V.; Scott, D. A.; Zhang, F., Genome engineering using the CRISPR-Cas9 system. Nat Protoc 2013, 8 (11), 2281–2308.

39. Hsu, P. D.; Scott, D. A.; Weinstein, J. A.; Ran, F. A.; Konermann, S.; Agarwala, V.; Li, Y.; Fine, E. J.; Wu, X.; Shalem, O.; Cradick, T. J.; Marraffini, L. A.; Bao, G.; Zhang, F., DNA targeting specificity of RNA-guided Cas9 nucleases. Nat Biotechnol 2013, 31 (9), 827–32.

40. Brinkman, E. K.; Chen, T.; Amendola, M.; van Steensel, B., Easy quantitative assessment of genome editing by sequence trace decomposition. Nucleic Acids Res 2014, 42 (22), e168.

41. Valgardsdottir, R.; Chiodi, I.; Giordano, M.; Cobianchi, F.; Riva, S.; Biamonti, G., Structural and functional characterization of noncoding repetitive RNAs transcribed in stressed human cells. Mol Biol Cell 2005, 16 (6), 2597–604.

42. Goenka, A.; Sengupta, S.; Pandey, R.; Parihar, R.; Mohanta, G. C.; Mukerji, M.; Ganesh, S., Human satellite-III non-coding RNAs modulate heat-shock-induced transcriptional repression. J Cell Sci 2016, 129 (19), 3541–3552.

43. Baum, P.; Fundel-Clemens, K.; Kreuz, S.; Kontermann, R. E.; Weith, A.; Mennerich, D.; Rippmann, J. F., Off-target analysis of control siRNA molecules reveals important differences in the cytokine profile and inflammation response of human fibroblasts. Oligonucleotides 2010, 20 (1), 17–26.

44. Winton, M. J.; Igaz, L. M.; Wong, M. M.; Kwong, L. K.; Trojanowski, J. Q.; Lee, V. M., Disturbance of nuclear and cytoplasmic TAR DNA-binding protein (TDP-43) induces disease-like redistribution, sequestration, and aggregate formation. J Biol Chem 2008, 283 (19), 13302–9.

45. Collins, M. A.; An, J.; Peller, D.; Bowser, R., Total protein is an effective loading control for cerebrospinal fluid western blots. J Neurosci Methods 2015, 251, 72–82.

46. Otter, T.; King, S. M.; Witman, G. B., A two-step procedure for efficient electrotransfer of both high-molecular-weight (greater than 400,000) and low-molecular-weight (less than 20,000) proteins. Anal Biochem 1987, 162 (2), 370–7.

47. Staudt, T.; Lang, M. C.; Medda, R.; Engelhardt, J.; Hell, S. W., 2,2’-thiodiethanol: a new water soluble mounting medium for high resolution optical microscopy. Microsc Res Tech 2007, 70 (1), 1–9.

48. Schneider, C. A.; Rasband, W. S.; Eliceiri, K. W., NIH Image to ImageJ: 25 years of image analysis. Nat Methods 2012, 9 (7), 671–5.

49. Rueden, C. T., B.; Schindelin, J.; Hiner, M., https://imagej.net/RATS.

50. Biamonti, G.; Vourc’h, C., Nuclear stress bodies. Cold Spring Harb Perspect Biol 2010, 2 (6), a000695.

51. Wang, X.; Zhou, S.; Ding, X.; Ma, M.; Zhang, J.; Zhou, Y.; Wu, E.; Teng, J., Activation of ER Stress and Autophagy Induced by TDP-43 A315T as Pathogenic Mechanism and the Corresponding Histological Changes in Skin as Potential Biomarker for ALS with the Mutation. Int J Biol Sci 2015, 11 (10), 1140–9.

52. Xu, Y. F.; Gendron, T. F.; Zhang, Y. J.; Lin, W. L.; D’Alton, S.; Sheng, H.; Casey, M. C.; Tong, J.; Knight, J.; Yu, X.; Rademakers, R.; Boylan, K.; Hutton, M.; McGowan, E.; Dickson, D. W.; Lewis, J.; Petrucelli, L., Wild-type human TDP-43 expression causes TDP-43 phosphorylation, mitochondrial aggregation, motor deficits, and early mortality in transgenic mice. J Neurosci 2010, 30 (32), 10851–9.

53. Alberti, S.; Mateju, D.; Mediani, L.; Carra, S., Granulostasis: Protein Quality Control of RNP Granules. Front Mol Neurosci 2017, 10, 84.

54. Lin, Y.; Protter, D. S.; Rosen, M. K.; Parker, R., Formation and Maturation of Phase-Separated Liquid Droplets by RNA-Binding Proteins. Mol Cell 2015, 60 (2), 208–19.

55. Uversky, V. N., Intrinsically disordered proteins in overcrowded milieu: Membrane-less organelles, phase separation, and intrinsic disorder. Curr Opin Struct Biol 2017, 44, 18–30.

56. King, O. D.; Gitler, A. D.; Shorter, J., The tip of the iceberg: RNA-binding proteins with prion-like domains in neurodegenerative disease. Brain Res 2012, 1462, 61–80.

57. Markmiller, S.; Soltanieh, S.; Server, K. L.; Mak, R.; Jin, W.; Fang, M. Y.; Luo, E. C.; Krach, F.; Yang, D.; Sen, A.; Fulzele, A.; Wozniak, J. M.; Gonzalez, D. J.; Kankel, M. W.; Gao, F. B.; Bennett, E. J.; Lecuyer, E.; Yeo, G. W., Context-Dependent and Disease-Specific Diversity in Protein Interactions within Stress Granules. Cell 2018, 172 (3), 590–604 e13.

58. Zhang, K.; Daigle, J. G.; Cunningham, K. M.; Coyne, A. N.; Ruan, K.; Grima, J. C.; Bowen, K. E.; Wadhwa, H.; Yang, P.; Rigo, F.; Taylor, J. P.; Gitler, A. D.; Rothstein, J. D.; Lloyd, T. E., Stress Granule Assembly Disrupts Nucleocytoplasmic Transport. Cell 2018, 173 (4), 958–971 e17.

59. Kim, H. J.; Taylor, J. P., Lost in Transportation: Nucleocytoplasmic Transport Defects in ALS and Other Neurodegenerative Diseases. Neuron 2017, 96 (2), 285–297.

60. James, B. D.; Wilson, R. S.; Boyle, P. A.; Trojanowski, J. Q.; Bennett, D. A.; Schneider, J. A., TDP-43 stage, mixed pathologies, and clinical Alzheimer’s-type dementia. Brain 2016, 139 (11), 2983–2993.

61. McAleese, K. E.; Walker, L.; Erskine, D.; Thomas, A. J.; McKeith, I. G.; Attems, J., TDP-43 pathology in Alzheimer’s disease, dementia with Lewy bodies and ageing. Brain Pathol 2017, 27 (4), 472–479.

62. Philips, T.; Rothstein, J. D., Glial cells in amyotrophic lateral sclerosis. Exp Neurol 2014, 262 *Pt B*, 111-20.

63. Boillee, S.; Vande Velde, C.; Cleveland, D. W., ALS: a disease of motor neurons and their nonneuronal neighbors. Neuron 2006, 52 (1), 39–59.

